# High-throughput determination of RNA tertiary contact thermodynamics by quantitative DMS chemical mapping

**DOI:** 10.1101/2024.03.11.584472

**Authors:** Bret Lange, Ricardo G. Gil, Joseph D. Yesselman

## Abstract

Structured RNAs often contain long-range tertiary contacts that are critical to their function. Despite the importance of tertiary contacts, methods to measure their thermodynamics are low throughput or require specialized instruments. Here, we introduce a new quantitative chemical mapping method (qDMS-MaPseq) to measure Mg^2+^-induced formation of tertiary contact thermodynamics in a high-throughput manner using standard biochemistry equipment. With qDMS-MaPseq, we measured the ΔG of 98 unique tetraloop/tetraloop receptor (TL/TLR) variants in a one-pot reaction. These results agree well with measurements from specialized instruments (R^2^=0.64). Furthermore, the DMS reactivity of the TL directly correlates to the stability of the contact (R^2^=0.68), the first direct evidence that DMS reactivity reports on thermodynamics. Combined with structure prediction, DMS reactivity allowed the development of experimentally accurate 3D models of TLR mutants. These results demonstrate that qDMS-MaPseq is broadly accessible, high-throughput, and directly links DMS reactivity to thermodynamics.

## Introduction

Structured RNAs play critical roles in cellular functions, are foundational to the life cycle of RNA viruses, and serve as blueprints for new artificial machines (1–6). RNAs must fold into intricate 3D structures to perform these diverse functions. This folding typically occurs in two distinct steps. Initially, nucleotide base pairs form, creating a secondary structure with helices, junctions, and loops. Subsequently, these secondary structures may engage in further long-range interactions - known as tertiary contacts - which exhibit a remarkable structure and stability range. For instance, some tertiary contacts are transient, involving single adenines docking into a helix’s minor groove (a-minor interactions) (7). In contrast, others encompass multiple base pairs or non-canonical interactions that provide enough thermodynamic stability to strongly favor a specific 3D conformation crucial for its function (8). Understanding the thermodynamics of these individual tertiary contacts is vital for unraveling how biological RNAs fold into their functional forms. Moreover, this knowledge will be instrumental in designing advanced artificial RNAs, opening new frontiers in biotechnology and medicine.

Due to the importance of tertiary contacts, several methods have been utilized to investigate their stability experimentally. Single-molecule FRET, native gel electrophoresis, small angle X-ray scattering, chemical probing, and others have all been used to assess tertiary contact stability quantitatively (6, 9–18). However, these time-intensive protocols require individual sample preparation, making them impractical for large-scale RNA studies. As the number of structured RNAs with potential tertiary contacts rapidly grows, there is a critical need for high-throughput methods. Greenleaf and colleagues have recently developed the RNA on a massively parallel array (RNA-MaP) platform (19–21). This device can simultaneously measure the stability of thousands of unique RNA tertiary contacts through fluorescence intensity. The RNA-MaP platform is a significant technological advance, leading to several important insights about tertiary contact formation (19–21). However, these devices are custom-built using existing Illumina sequencers and, thus, are not accessible to other scientists. Consequently, the scientific community still faces an urgent need for a simple, high-throughput method that is widely accessible.

Chemical modifiers, such as dimethyl sulfate (DMS), offer an alternative for measuring tertiary contact formation in RNAs. DMS methylates the N1 position of adenine and the N3 position of cytosine when accessible to solvent. These DMS reactivity profiles are traditionally employed to constrain RNA secondary structures, as the targeted nitrogen in Watson-Crick pairs is shielded from the DMS-induced methylation (22–26). However, several tertiary contacts, including kissing loops, pseudoknots, A-minor interactions, and the tetraloop/tetraloop receptor (TL/TLR), also protect against DMS methylation (6, 27–30). Recently, DMS has been coupled with next-generation sequencing techniques such as mutational profiling with sequencing (DMS-MaPseq) (24–26). DMS-MaPseq allows for simple multiplexing of the DMS reaction to probe the DMS reactivity of thousands of RNAs in a one-pot reaction. DMS-MaPseq has led to multiple advances in understanding conformational changes in secondary structure but has yet to be used to measure 3D structure thermodynamics.

In physiological environments, the formation of many tertiary RNA contacts relies on divalent cations, particularly Mg^2+^, to mitigate the electrostatic repulsion of the phosphate backbone. Decades of research have demonstrated that the concentration of Mg^2+^ required for a tertiary contact to form is directly related to its thermodynamic stability. The concentration of Mg^2+^ required for 50% of the RNA molecules to form their tertiary contact is known as the [Mg^2+^]_½_. A low [Mg^2+^]_½_ requirement suggests a more stable interaction capable of overcoming inherent electrostatic repulsion. Furthermore, using older reverse transcription stop methods, DMS chemical mapping has been employed to measure [Mg^2+^]_½_ for individual RNA constructs (6). We posited that DMS-MaPseq might allow simultaneous measurement of [Mg^2+^]_½_ for hundreds of RNAs in one-pot reactions. This new method would bring high-quality thermodynamic measurements to a much more extensive range of researchers.

This study introduces a new quantitative DMS-MaPseq method (qDMS-MaPseq) that integrates [Mg^2+^] titrations to measure the thermostability of tertiary contacts. qDMS-MaPseq can be performed on 1 to over 10,000 RNAs in a one-pot reaction, provides nucleotide-resolution data, is low cost, and uses common reagents found in any molecular biology lab. To benchmark qDMS-MaPseq, we used an RNA nanostructure containing a tetraloop/tetraloop receptor (TL/TLR) tertiary contact. We measured the thermostability of 98 TLR mutants and obtained results that were consistent with the specialized RNA-MaP platform. Surprisingly, the DMS reactivity of the TL directly correlates to the stability of the contact (R^2^=0.68), the first direct evidence that DMS reactivity reports on thermodynamics. qDMS-MaPseq data yields rich nucleotide-level data that contains embedded information about 3D structures. These data permitted observation of an Mg^2+^-induced conformational change in the TLR that explains discrepancies between the free TLR and TLR-bound structures. Lastly, we used these data to build preliminary 3D models of TLR mutant structures. Our findings indicate that qDMS-MaPseq can democratize and generalize high-quality thermodynamic methods of tertiary contacts that can be applied to a wide range of systems.

## Results

### Developing a quantitative DMS-MaPseq protocol to observe Mg^2+^-induced tertiary contact formation

To probe the thermodynamics of tertiary contact formation, we utilized a minimal tetraloop/tetraloop receptor (miniTTR) nanostructure as a model system (**Figure 1A**). We used DMS-MaPseq to probe the structure of the miniTTR construct (**Figure 1B**) (24–26). The tetraloop and tetraloop-receptor have near-zero mutation fraction (DMS reactivity) in 10 mM Mg^2+^, indicating that contact formation shields the N1 position of the four interacting adenines from methylation, consistent with previous DMS probing results (**Figure 1B, purple circles**) (6). To observe the Mg^2+^-induced formation of the TL/TLR, we performed the standard DMS-MaPseq protocol with and without 10 mM Mg^2+^. We observed no significant difference in reactivity between the conditions (**Figure 2A**). We postulated that the ionic strength (300 mM sodium cacodylate) in the standard DMS-MaPseq experiment enabled sodium to provide sufficient electrostatic screening without divalent counter ions, a previously observed effect (31, 32).

**Figure 1:**
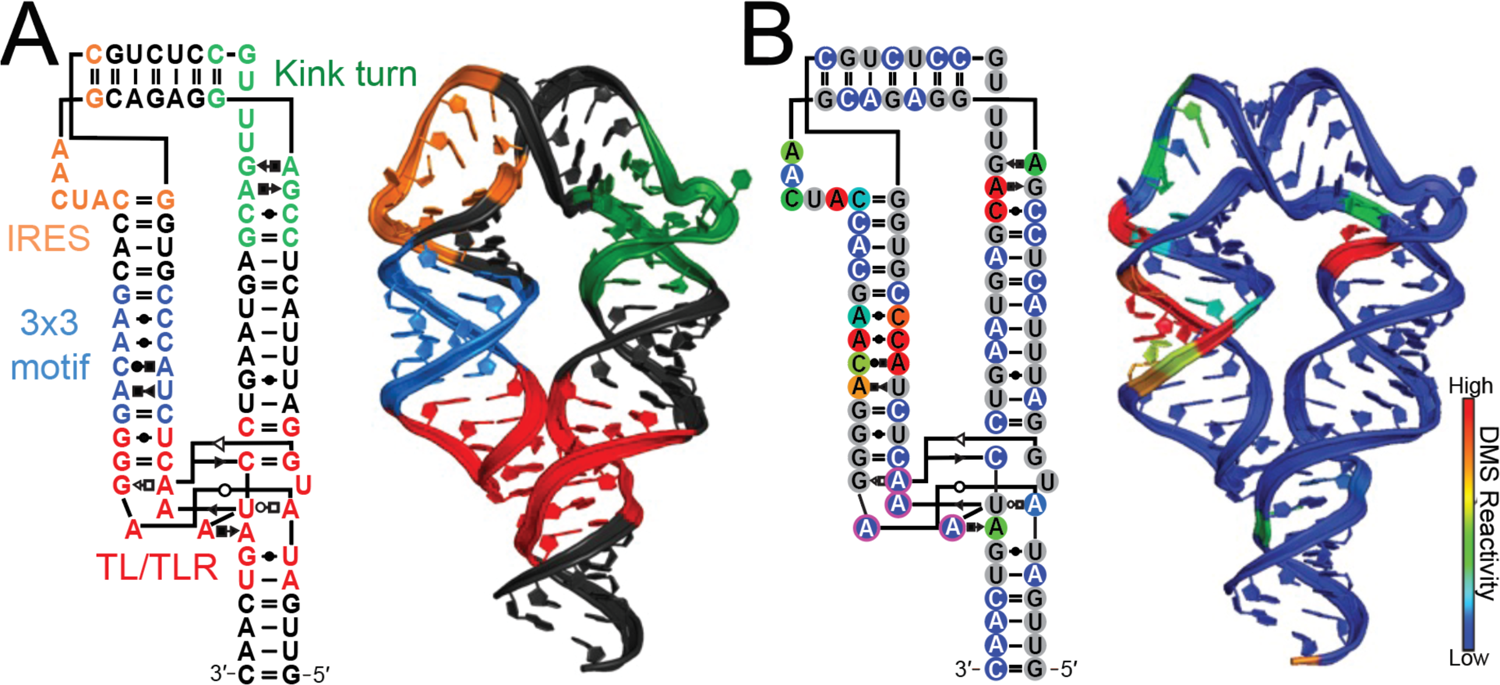
Overview of minimal tetraloop/tetraloop receptor construct. (A) The secondary structure and tertiary structure (PDB: 6DVK) are colored by secondary structure motifs, helices are in black. (B) The secondary and tertiary structures are colored by DMS reactivity, indicating the tetraloop/tetraloop receptor is formed. The 4 adenines that can serve as probes of tertiary contact formation are highlighted in magenta.

**Figure 2:**
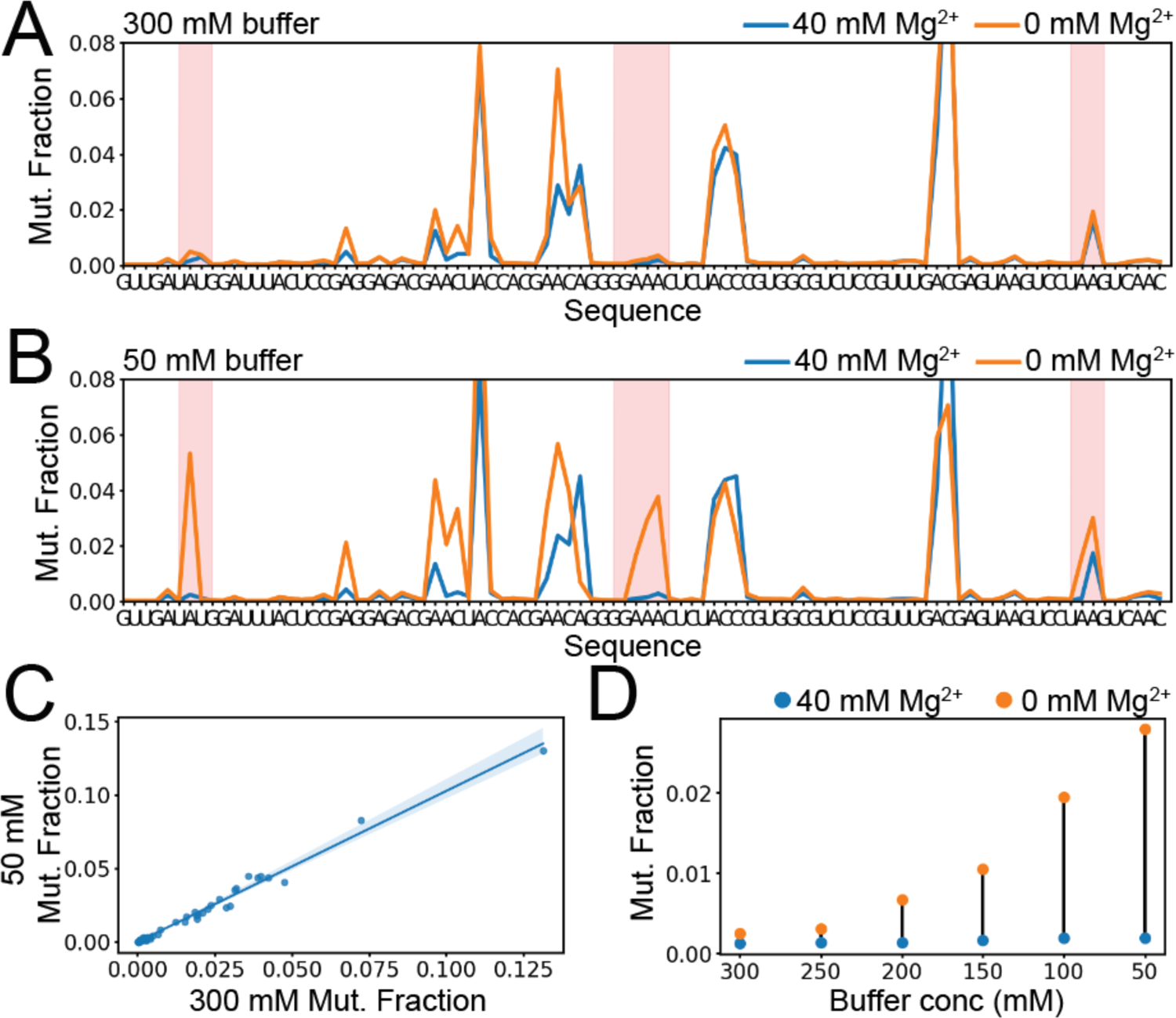
Buffer effects on the DMS reaction and tertiary contact stability. (A) Using 300 mM sodium cacodylate, compare 40 mM Mg^2+^ to 0 mM Mg^2+^ with no notable difference in the TL/TLR mutational fraction. (B) A comparison between 40 mM Mg^2+^ and 0 mM Mg^2+^ with 50 mM sodium cacodylate showed notable differences between the mutation fraction in the TL/TLR (highlighted in red), indicating tertiary contact formation. (C) The high correlation between DMS mutation fraction at 50 mM and 300 mM sodium cacodylate at 10 mM Mg^2+^ indicates that the lower buffer is sufficient to buffer the reaction and gives near identical data. (D) The change in mutation fraction of the GAAA tetraloop between 0 and 40 mM Mg^2+^ displaying increasing changes as the buffer concentration increases.

To determine the effect of [Na^+^] on the reactivity of the miniTTR construct, we performed DMS-MaPseq at six sodium cacodylate concentrations (300, 250, 200, 150, 100, and 50 mM) (**Figure 2, Supplemental Figure 1**) with and without 40 mM Mg^2+^. The reactivity difference in the TL/TLR between the 0 and 40 mM Mg^2+^ increased as the amount of [Na^+^] decreased. The 50 mM sodium cacodylate condition demonstrated the largest effect (**Figure 2D**). Moreover, the 0 mM Mg^2+^ condition mirrors the reactivity profile of two negative control constructs that cannot form the tertiary contact (**Supplemental Figure 2-3**). Lastly, a high correlation (R^2^ = 0.96) was observed between the reactivity of the 300 mM and 50 mM sodium cacodylate conditions with 40 mM Mg^2+^ (**Figure 2C**). This correlation indicates that the decrease in sodium cacodylate does not cause observable changes in structure. Thus, we concluded that 50 mM sodium cacodylate is the optimal buffer condition for observing Mg^2+^-induced tertiary contact formation by DMS-MaPseq.

**Figure 3:**
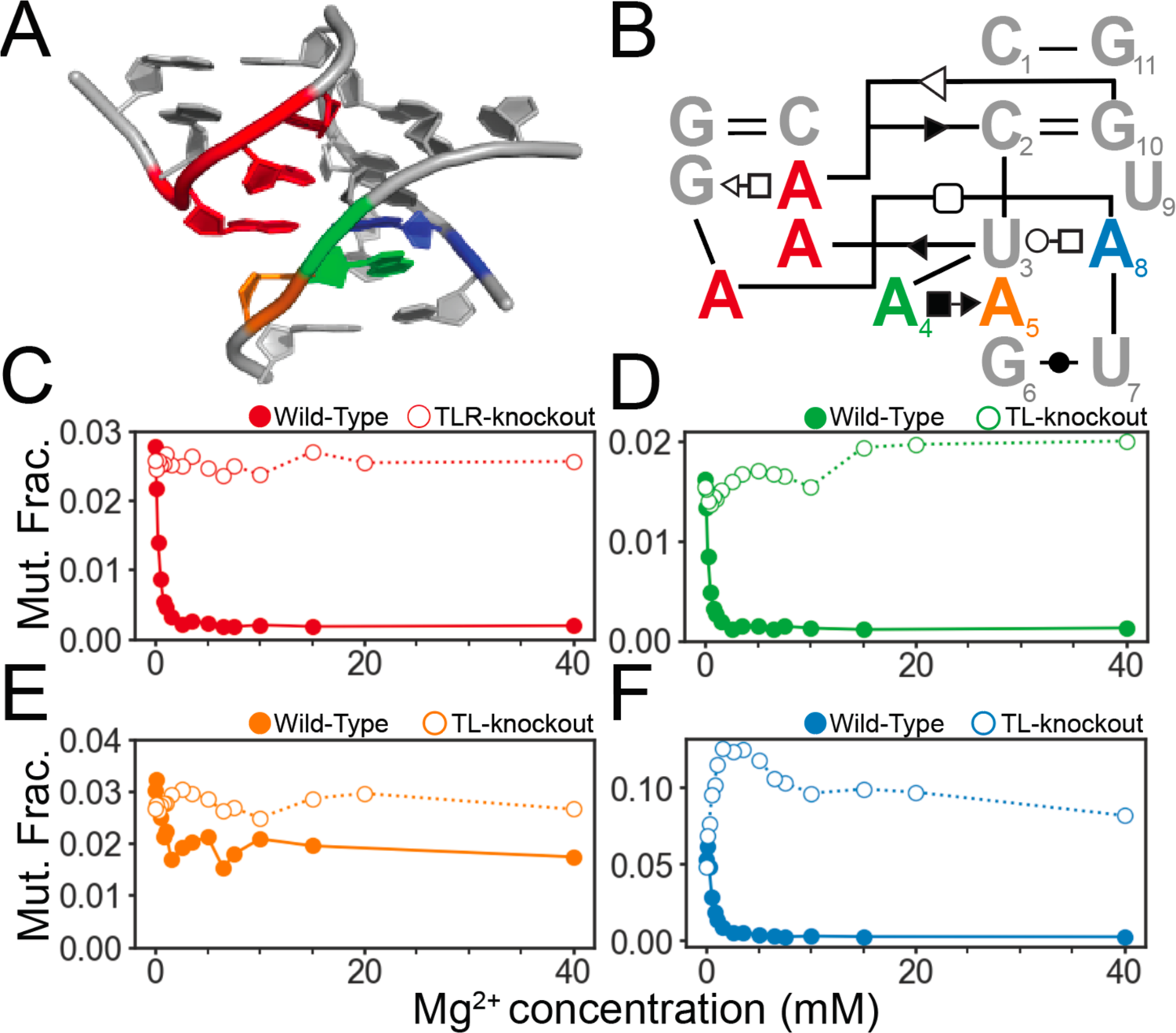
Nucleotide level conformational information as a function of [Mg^2+^]. (A) 3D structure of the TL/TLR interaction. Colored by residue, red is the three adenines in the tetraloop, green is A4, orange is A5, and blue is A8 of the tetraloop receptor. (A) the 3D structure of the TL/TLR interaction. (B) The secondary structure of the TL/TLR interaction. (C) The mutation fraction of the average of the three adenines of tetraloop in filled circles. The mutation fraction of a TLR-knockout in open circles is missing the TLR (**Supplemental** Figure 7) and thus cannot form the contact. (D) The mutation fraction of A4 of the TLR in the wild-type (closed circles) and the TL-knockout (open circles). (E) The mutation fraction of A5 of the TLR in the wild-type (closed circles) and the TL-knockout (open circles). (F) The mutation fraction of A8 of the TLR in the wild-type (closed circles) and the TL-knockout (open circles).

### DMS-MaPseq interrogates the behavior of individual nucleotides during TL/TLR formation

We performed DMS-MaPseq at 16 Mg^2+^ concentrations to investigate how each adenine’s DMS reactivity changes in the TL/TLR contact of the miniTTR scaffold. We engineered two control variants to isolate reactivity changes attributable to tertiary contact formation. The first control variant was a TL-knockout, which substitutes the GAAA tetraloop with a UUCG incapable of forming the TL/TLR contact. The second control was a TLR-knockout, where the TLR is replaced by a helix (**Supplemental Figure 2**). First, we examined the reactivity change as a function of Mg^2+^ concentration for the three adenines in the tetraloop (**Figure 3B**). All three tetraloop adenines behave similarly, so we averaged their reactivity (**Supplemental Figure 4**). We observe a rapid decrease in their average reactivity approaching 0 as Mg^2+^ concentration increases (**Figure 3C, closed circles**), indicative of TL/TLR formation. In contrast, the reactivity of tetraloop adenines within the TLR-knockout remains constant as Mg^2+^ concentration increases (**Figure 3C, open circles**).

Next, we analyzed the reactivity trends for each adenine in the tetraloop-receptor as a function of Mg^2+^ concentration. A8 forms an A-A pair with the tetraloop that involves its N1 position; thus, it behaves similarly to the tetraloop’s three adenines with a rapid reactivity decrease as Mg^2+^ concentration increases (**Figure 3F, closed circles**). However, in the TL-knockout, A8 reactivity rapidly increases, followed by a plateau as a Mg^2+^ concentration increases. (**Figure 3F, open circles**). The rapid increase in reactivity suggests a conformational change induced by Mg^2+^. We observed a similar behavior of the A4 reactivity profile of the TLR as a function of Mg^2+^ concentration. In the presence of the GAAA, A4’s reactivity rapidly decreases, and in the TL-knockout it increases and then plateaus. A4 does not form direct contact with the tetraloop but does have N1 positions facing out into the solvent if it retains the same conformation without the tetraloop present (**Figure 3D**). A5, which forms the A platform under A8, undergoes a slight decrease in reactivity as a function of Mg^2+^ concentration with the GAAA tetraloop present but not to the same magnitude as A4 and A8. Furthermore, A5’s reactivity does not sharply increase with Mg^2+^ concentration with the TL-knockout, suggesting its N1 atom’s local environment does not significantly change with Mg^2+^ concentration.

### Tetraloop-receptor nucleotide reactivity profiles indicate Mg^2+^-induced conformational change independent of tetraloop binding

Based on the rapid increases in reactivity of A4 and A8 as a function of Mg^2+^ concentration in the TL-knockout, we hypothesized that the TLR undergoes a Mg^2+^-induced conformational change independent of tetraloop binding. To investigate this potential conformational change, we examined the NMR structure of the tetraloop receptor in the absence of its tetraloop binding partner (PDB: 1TLR) (33). This structure was resolved without Mg^2+^ and does not contain the iconic A-A platform between A4 and A5 (**Figure 4A**). This is consistent with previous research demonstrating that the A-A platform does not form in low ionic strength without divalent ions (34, 35). One possible explanation for the rapid increase of reactivity of A4 and A8 is that upon adding Mg^2+^, we are observing the formation of an A-A platform, which forces both A4 and A8 into conformations that are much more solvent exposed.

**Figure 4:**
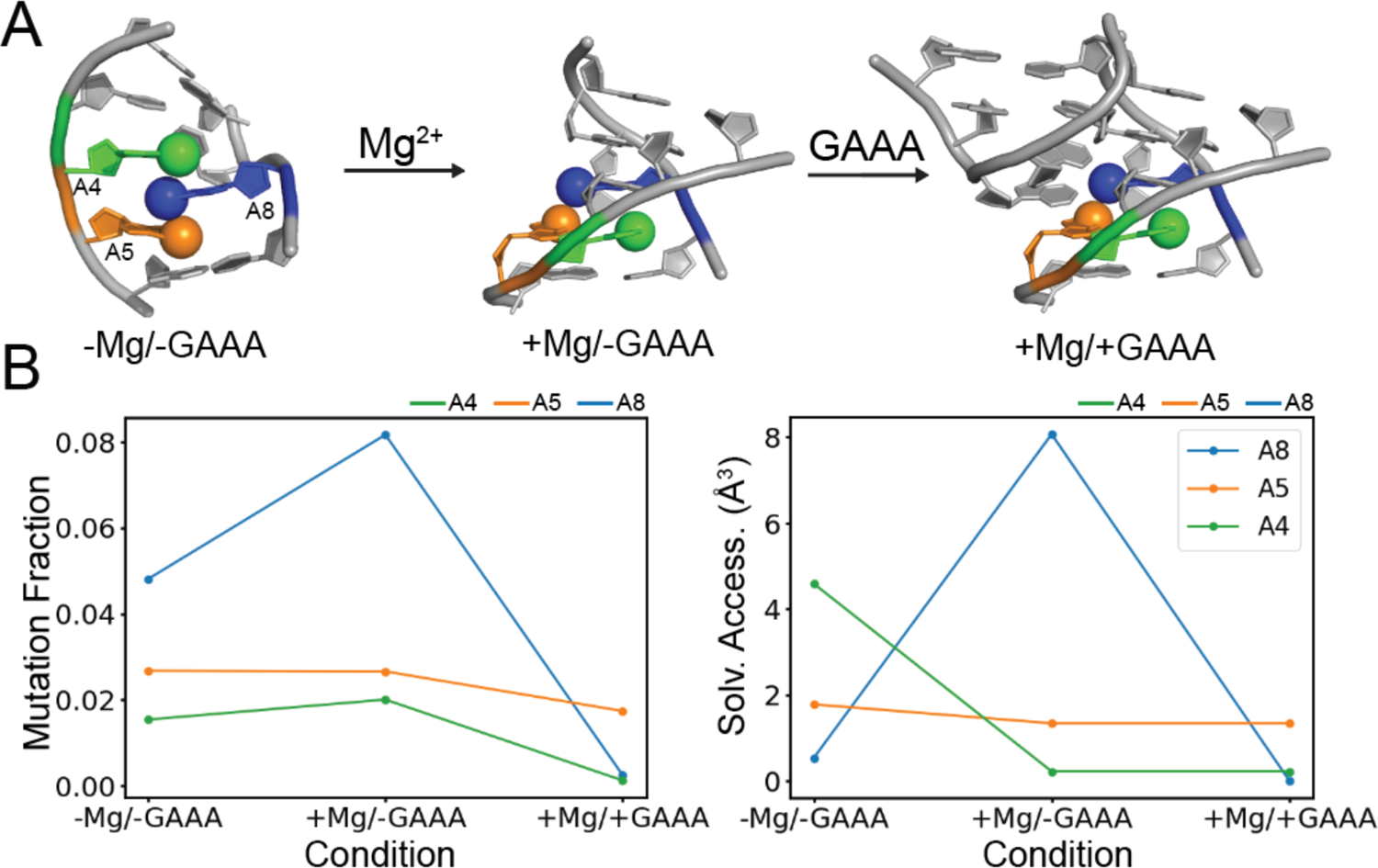
DMS mutation fractions at the postulated three states of the tetraloop receptor. -Mg/- GAAA state is without Mg^2+^ and GAAA tetraloop (PDB: 1TLR). +Mg/-GAAA state is with Mg^2+^ but no GAAA tetraloop is present (PDB: 1GID, tetraloop removed). +Mg/+GAAA state has both Mg^2+^ and GAAA tetraloop (PDB: 1GID). (A) The proposed three state model of the TLR, A4 (green), A5 (orange), A8 (blue). (B) On the left, the mutation fraction of A4, A5 and A8 over the three states. On the right the solvent accessibility of the N1 position of A4, A5, A8 over the three states.

To evaluate this hypothesis, we reasoned that our DMS reactivity should be correlated to the solvent accessibility for each conformation. To compute the solvent accessibility, we used the PDBs 1LTR for the no Mg^2+^ and no GAAA tetraloop state (-Mg/-GAAA) and 1GID for both the +Mg/-GAAA (removing the tetraloop) and +Mg/+GAAA state. A8 change in solvent accessibility over the three states mirrors its DMS reactivity (**Figure 4B-C**). A5 maintains consistent solvent accessibility across all three states, with a corresponding stable DMS reactivity trend. A4 presents a slight outlier case; its solvent accessibility is high in the -Mg/-GAAA state but decreases substantially in both the +Mg/+GAAA states, only partially aligning with its DMS reactivity. It may be that A4 has a broader set of conformations that are not fully explained by the apo TLR structure. These data are consistent with an A-A platform upon adding Mg^2+^ ions but are not definitive and require additional experiments.

### Measuring the effects of structural mutations destabilizing the TL/TLR interaction using qDMS-MaPseq

To measure the stability of the miniTTR TL/TLR tertiary contact, we measured the [Mg^2+^]_½_ value of 0.22 ± 0.004 mM from our 16-point Mg^2+^ titration. We denote using DMS-MaPseq to measure thermodynamic parameters as quantitative DMS-MaPseq or (qDMS-MaPseq). This value is similar to our previous study with different buffer conditions (0.12 ± 0.03 mM) (6). To assess the sensitivity of qDMS-MaPseq, we generated three destabilizing mutations, each designed by extending one of the helices linking the tetraloop and its receptor (**Figure 5, Supplemental Figure 6**). We hypothesize that these helix insertions will disrupt the precise alignment necessary for docking between the tetraloop and tetraloop-receptor. We performed a 16-point Mg^2+^ titration with qDMS-MaPseq and determined observed a significantly higher [Mg^2+^]_½_ for each mutant of 1.10 ± 0.25 mM, 1.37 ± 0.17 mM, and 0.58 ± mM for the helix H1, H2, and H3 insertions, respectively (**Supplemental Figure 7-9**). These elevated [Mg^2+^]_½_ values indicate a reduced frequency of tertiary contact formation in the mutants compared to the wild-type scaffold. The wild-type construct has an average GAAA reactivity of 0.002 at 40 mM Mg^2+^. The average GAAA reactivity at 40 mM Mg^2+^ increases for each destabilized mutant and is linearly correlated with their [Mg^2+^]_½_ values (**Supplemental Figure 10**). This indicates that at the 40 mM Mg^2+^, these mutants have their tertiary contact formed less frequently than the wild-type.

**Figure 5:**
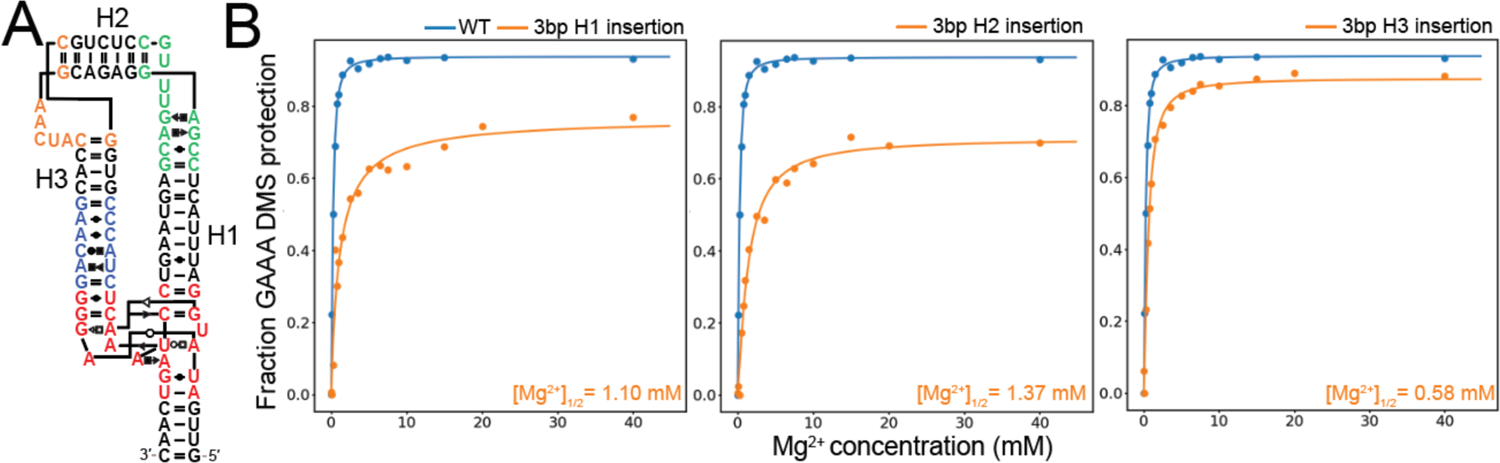
qDMS-MaPseq can measure the destabilization of TL/TLR due to helix lengthening. (A) For a schematic of which helices will be lengthened, see **Supplemental** Figure 11 for secondary structures of each insertion construct compared to the wild-type. (B) Left is a comparison between the mutation fraction of the wild-type and the 3-bp helix 1 (H1) insertion construct. Middle, a comparison between the mutation fraction of the wild-type and the 3-bp helix 2 (H2) insertion construct. Right, a comparison between the mutation fraction of the wild-type and the 3-bp helix 3 (H3) insertion construct.

An intriguing aspect of these results is the asymmetrical impact of the mutations. Insertions in helices H1 and H2 led to more pronounced destabilization than H3. Converting their [Mg^2+^]_½_ to ΔΔG with the wild-type as a reference has H2 destabilized by 1.13 kcal/mol compared to H3 of 0.54 kcal/mol. This suggests that insertions into H3 are more easily compensated, potentially by breaking the weak non-canonical interactions in the 3×3 motif (6). These results indicate that qDMS-MaPseq can measure small destabilizations throughout the miniTTR scaffold, probing how well the TL/TLR contact is aligned.

### qDMS-MaPseq can measure the ΔG of TL/TLR mutants in a high throughput manner

In a recent set of studies, Herschlag and colleagues characterized the binding affinities of approximately 3000 tetraloop receptor mutants to the GAAA tetraloop using the RNA-MaP platform (19, 21). They identified a subset of mutations with the same binding mode as the wild-type tetraloop-receptor (11ntR). This subset presents a unique opportunity to compare the [Mg^2+^]_½_ of qDMS-MaPseq to existing thermodynamic measurements. We selected a representative sample of 98 mutants from this subset for our initial proof-of-concept experiment. The range of these binding affinities in this set to the GAAA tetraloop receptor ranges from −11.77 to −8.35 kcal/mol (**Supplemental Table 1**). Each mutant sequence was incorporated into the miniTTR scaffold, replacing the standard 11ntR receptor. Our naming convention uses the TLR mutant sequence. For example, a mutation such as A2C has a sequence (C**A**UAA&UAUGG); thus, we name the miniTTR construct with this mutated TLR: C**A**UAA_UAUGG (**Figure 6A**). We performed our standard 16-point [Mg^2+^] titration protocol on this set of mutated TLR constructs. We determined the [Mg^2+^]_½_ values for 84 of the 98 mutants. Of the remaining 14, 11 were below the detection range of qDMS-MaPseq, exhibiting [Mg^2+^]_½_ values well beyond our testing range of 40 mM. These mutants corresponded to the lowest ΔG values in our dataset, ranging from −9.50 to −8.30 kcal/mol, as outlined in **Supplemental Table 1**. The other three constructs excluded were for issues with the DMS reactivity. In UCUAAA_CAUGA and CCUACA_UACGG, we observed significant reactivity in the G=C closing pair to the GAAA tetraloop, affecting its binding behavior (**Supplemental Figure 11**). Finally, in CUUAAC_UAUGG, its GAAA tetraloop did not reduce reactivity as expected but changed into a new unknown reactivity pattern, suggesting an alternative structure (**Supplemental Figure 12**). Our analysis revealed a strong correlation between the natural log of [Mg^2+^]_½_ values obtained from qDMS-MaPseq and the ΔG values from RNA-MaP, achieving an R^2^ of 0.64 (**Figure 6C**). These results indicate that qDMS-MaPseq produces comparable thermodynamic measurements to the established RNA-MaP experiments.

**Figure 6:**
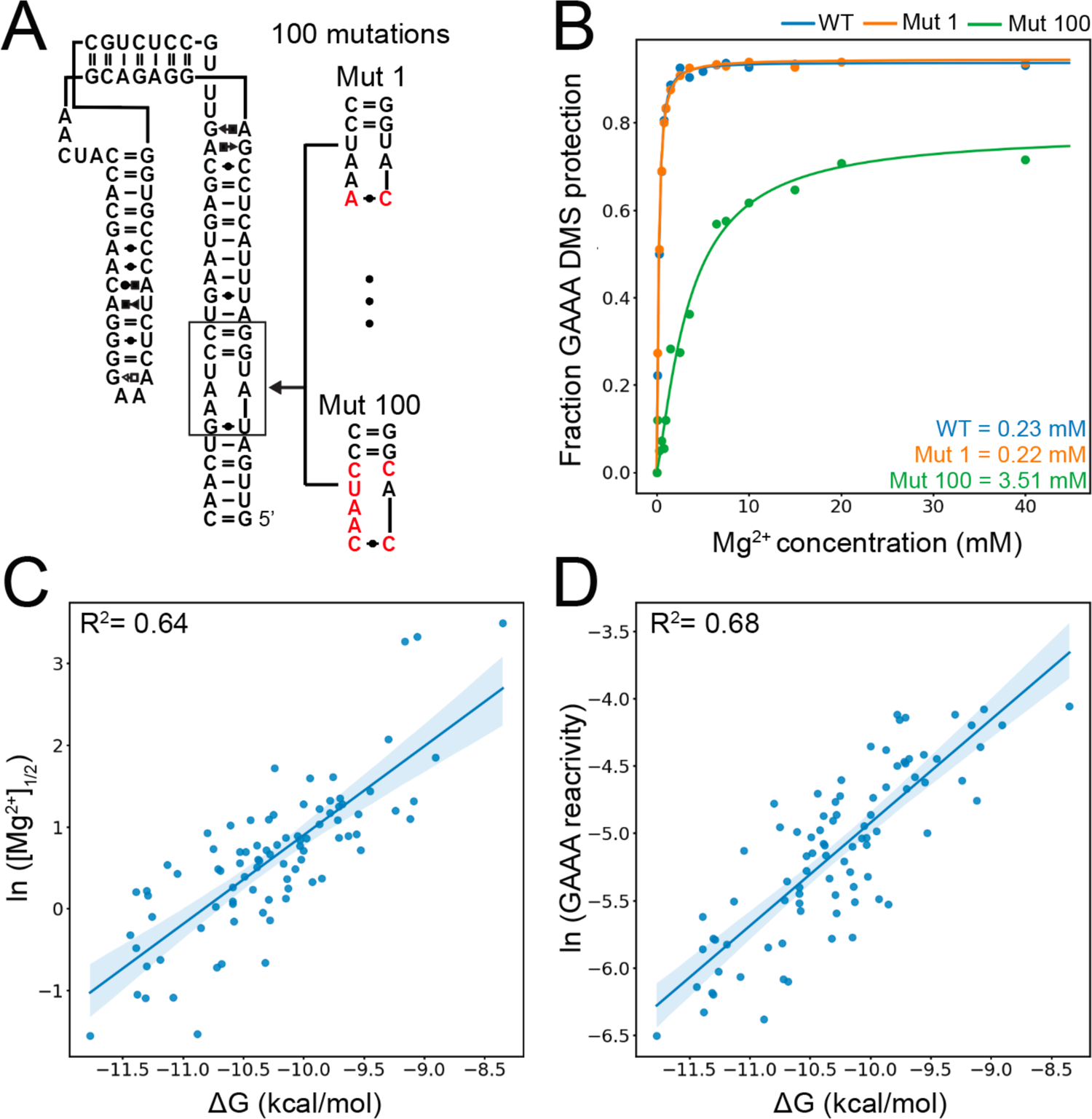
Comparison between qDMS-MaPseq results and known ΔG values for 98 TLR mutations. (A) A schematic of how mutant TLRs were inserted into the miniTTR scaffold. (B) Comparison between Mg^2+^ titrations of the wild-type and two TLR mutants. (C) Correlation plot between the natural log of [Mg^2+^]_½_ and the ΔG measured by the RNA-MaP experiments from Bonilla et al. for each TLR mutation. (D) Correlation plot between the natural log of average DMS reactivity of the three adenines compared to the ΔG measured by the RNA-MaP experiments from Bonilla et al. for each TLR mutation.

We further noted that the average reactivity of the three As in the GAAA tetraloop at multiple [Mg^2+^] concentrations was correlated with the [Mg^2+^]_½_ values. For example, at 7.5 mM Mg^2+^, simply taking the average reactivity of the three As in the GAAA tetraloop gives a coefficient of determination of 0.77 compared to the [Mg^2+^]_½_ values (**Supplemental Figure 13**). Thus, we compared the natural log of the average reactivity of three adenines in the GAAA tetraloop at 7.5 mM [Mg^2+^] to the ΔG measured by RNA-MaP (Figure 6D). Surprisingly, we achieved a higher correlation coefficient of R^2^ = 0.68. This suggests that DMS reactivity values contain significant information content and are directly correlated to thermodynamic parameters. This correlation does not only exist at 7.5 mM Mg^2+^ to the ΔG but at many unique Mg^2+^ concentrations (**Supplemental Figure 14**). These results underscore the predictive power of a single DMS reactivity measurement, simplifying the experimental protocol and enhancing its accessibility.

### qDMS-MaPseq identifies residue-specific structural features on tetraloop receptor mutants

In addition to the accessibility and simplicity, qDMS-MaPseq offers nucleotide-level insights into how mutations alter the 3D structure of the tetraloop receptor. We examined three repeating TLR mutations that yielded unique DMS reactivity profiles. For these TLR mutants, we generated potential 3D models utilizing Fragment Assembly of RNA with Full Atom Refinement (FARFAR) (36, 37). Previously, we used FARFAR to build a 3D model of the 12-ntR receptor that could successfully explain the effects of known mutations (20, 38).

First, we examined a G5C/U6C variant, which mutates the G-U pair to a C-C pair (**Figure 7A**). This mutation occurs 31 times (**Supplemental Table 2**). CCUAA**C**_**C**AUGG is the only variant that contains this G-U to C-C mutation. This mutation produces a striking reactivity pattern change with and without 40 mM Mg^2+^. At 0 mM Mg^2+^, C5 and C6 have high and comparable reactivities of 0.021 and 0.032, respectively. However, at 40 mM Mg^2+^, C5’s reactivity drops to 0.009, and C6’s reactivity increases to 0.064. C6’s reactivity is 7-fold higher than C5’s, indicating a significant difference in the chemical environments of their N1 atoms (**Figure 7A**). To understand this radial asymmetry in reactivity in this C-C, we examined the ten lowest energy structures generated by FARAFAR (**Figure 7B**). These models converged on a potential explanation. C5’s N1 forms a hydrogen bond to one of the hydrogens on C6’s N4 (**Figure 7C**). This hydrogen bond likely shields C5’s N1 from methylation and holds C6’s N1 pointed out into solvent, leading to higher methylation levels. This unique asymmetric C-C pair is also in the miniTTR scaffold’s kink-turn and yields a similar reactivity profile (**Figure 7D and Supplemental Figure 15**).

**Figure 7:**
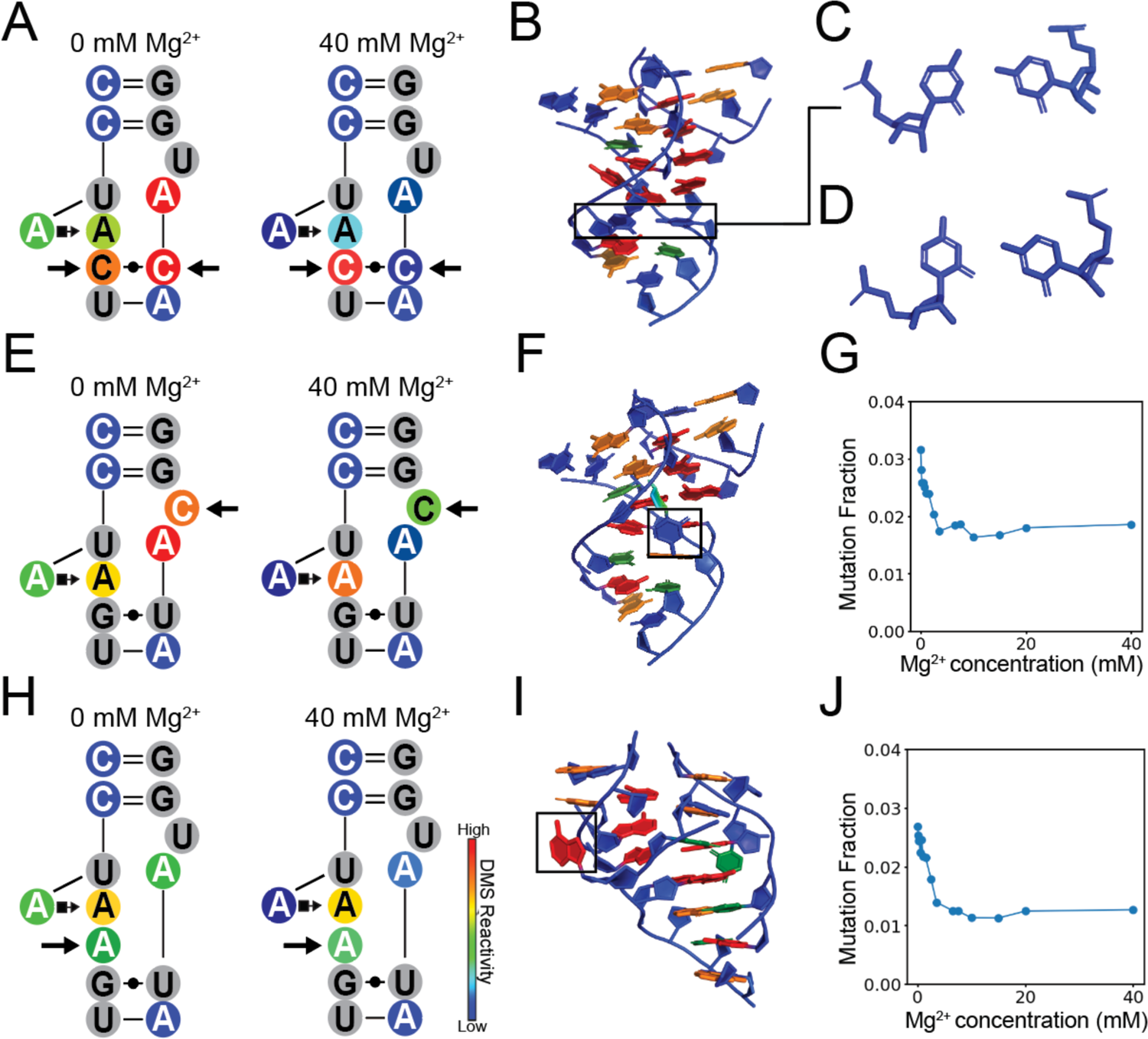
DMS reactivity signatures indicate 3D structure features in TLR mutants. (A) DMS reactivity of CCUAAC_CAUGG with and without 40 mM Mg^2+^ displaying the unique DMS signature of the C-C mutation highlighted with arrows. (B) FARFAR generated model of CCUAAC_CAUGG TLR mutant, boxed is the C-C mismatch. (C) The FARFAR model of the C-C mismatch, which has a unique asymmetrical orientation, with the right C turned to form a hydrogen bond with the N1 position of the left C. (D) The C-C mismatch from the kink-turn motif in the miniTTR scaffold (PDB: 6DVK) has a unique asymmetrical orientation, with the right C turned to form a hydrogen bond with the N1 position of the left C. This asymmetry is likely the cause of its skewed DMS reactivity, where the right C is over 10-fold more reactive than the left C. (E) DMS reactivity of CCUAAG_UACGG with and without 40 mM Mg^2+^ with a C9U mutation. (F) In the FARAFAR model of the C9U mutation, this C is flipped out like the wild-type U. (G) The DMS reactivity as a function of Mg^2+^ where the C levels out to reactivity indicating is consistently solvent accessible. (H) DMS reactivity of CCUAAAG_UAUGG with and without 40 mM Mg^2+^ with an A insertion. (I) In the FARAFAR model of the CCUAAAG_UAUGG, the inserted A is flipped out and does not participate in the A-A platform. (J) The DMS reactivity as a function of Mg^2+^ where the inserted A levels out to reactivity, indicating that it is consistently solvent accessible and has a unique profile that differs from the As in the A-A platform.

Next, we examined a C9U mutation. **Figure 7E**). This U is flipped out and does not form any contact with the tetraloop in the crystal structure of the TL/TLR complex. We examined the construct CCUAAG_UA**C**GG, which contains only this C9U mutation, where we hypothesize the C will also be flipped out. The reactivity at this C decreases rapidly at low [Mg^2+^] levels, consistent with an initial conformation change induced by the Mg^2+^ (**Figure 7J**). The top 10 models converge highly for this sequence, and all have C9 flipped out — the reactivity at higher Mg^2+^ levels is about 0.015, similar to other flipped Cs (**Supplemental Figure 16**).

Finally, we investigated an intriguing insertion mutation, where an additional adenine is added at position 4, resulting in a sequence of three consecutive adenines (CCU**A**AAG_UAUGG). This modification poses questions about the role of the newly inserted A in the A-A platform formation. Comparing the reactivity of A4 and A5 in this mutant with the wild-type shows similar behaviors, suggesting they perform analogous roles. However, the inserted A exhibits a unique reactivity profile (**Figure 7J**), not aligning entirely with the behavior of A4 or A5. This reactivity pattern resembles the C9U mutation in CCUAAG_UA**C**GG, leading us to hypothesize that this A is flipped out and does not directly participate in tetraloop binding. One of the FARFAR models supports this hypothesis (**Figure 7I**). However, it should be noted that there are lower energy models that have A5 flipped out, but that does not agree with our experimental data. These findings indicate that, unlike previously high-throughput methods, the nucleotide-level data generated by qDMS-MaPseq can provide details into RNA’s 3D structure.

## Discussion

In this study, we introduced qDMS-MaPseq, a novel chemical mapping technique designed to interrogate tertiary contact in a rapid, high-throughput fashion. A significant advantage of qDMS-MaPseq over prior methods is its accessibility: it does not require specialized equipment, extensive training, or sequence design. This makes it highly applicable to various tertiary contact systems that depend on Mg^2+^, such as pseudoknots and kissing loops, vital in the genomes of viruses like HIV-1 and SARS-CoV-2 (24, 30). DMS-MaPseq has been demonstrated to work on complete viral genomes, eliminating size as a potential hindrance. Furthermore, there is no limit on how many sequences can be done in a single library. If it’s possible to differentiate each unique sequence in principle, one can examine millions of unique sequences in a single reaction. This is achievable with current technology, as oligo pools of this size are commercially available, and sequencing continues to decrease in price. This scalability, coupled with decreasing sequencing costs and the burgeoning field of machine learning, opens up exciting possibilities for exploring the fundamental rules of tertiary contact formation at an unprecedented scale.

Our observation that the natural logarithm of the GAAA tetraloop’s reactivity correlates more closely with the ΔG of TLR mutants than the [Mg^2+^]_½_ was unexpected. This finding simplifies the data collection, requiring fewer data points and revealing that DMS reactivity values contain more nuanced information than the traditional binary high/low reactivity interpretation. This marks the first time raw DMS reactivity has been directly correlated with structural thermodynamics beyond the Watson-Crick base pairing. Previous research has indicated that high reactivity often signals a contact break, and low reactivity indicates formation (39–41). Our findings add a new quantitative fundamental (ΔG) dimension. This methodology is readily extended to other non-canonical interactions.

In summary, qDMS-MaPseq emerges as a powerful tool for probing the 3D structure and stability of RNA, paving the way for future studies in RNA biology. Its capability to provide detailed, high-throughput analysis holds significant promise for advancing our understanding of RNA structure-function dynamics and potentially contributing to developing new therapeutic strategies targeting RNA in various diseases.

### Data, materials, and software availability

All data, materials, and software used in this study are available. Unprocessed FASTA files have been deposited to the Sequence Read Archive (SRA) under the accession PRJNA1086549. All other data is available on Fig Share (http://doi.org/10.6084/m9.figshare.25331758). All code used in this study is available on GitHub (https://github.com/jyesselm/q_dms_ttr_paper)

## Acknowledgments

This work was supported by the NSF (NSF CAREER 214363) to J.D.Y. We would like to thank Catherine Eichhorn, Daniel Herschlag, and Silvi Rouskin for their thoughtful comments, which strengthened this paper.

## Contributions

J.D.Y designed the experiments and analyzed the data. B.L. and R.G.G. performed the experiments. J.D.Y. wrote the paper with the help of B.L. and R.G.G.

## Methods

### Designing TLR mutant libraries

Using the supplemental data from Bonilla et al. (19), we selected a test set from that study’s original 1592 TLR constructs. First, we only considered mutants that fell into class A (277), which had a thermodynamic fingerprint similar to the wild-type. We reasoned that we only wanted to consider mutant binding modes similar to the wild-type for this first proof-of-concept study. We further filtered out constructs with few A and C nucleotides, removing all with less than three total A and C nucleotides. Lastly, we removed constructs that did not contain two Watson-Crick pairs flanking the tetraloop receptor. This left us with 98 mutants. We broke these into four sets that either had 25 or 24 constructs. We randomly selected helical sequences to ensure distinguishability among these mutants, as depicted in **Supplemental Figure 17**. Initial studies indicate that changing the helical sequence does not alter the reactivity of the tetraloop or receptor. Furthermore, we extended the first helix by seven base pairs to include a unique barcode preceding the tetraloop receptor, achieving a hamming distance of 20 or greater between each sequence.

### Ordering primers and oPools from Integrated DNA Technologies (IDT)

For the wild-type miniTTR 6, miniTTR 6 TL-knockout, miniTTR 6 TLR-knockout, miniTTR 6 h1 3bp insert, miniTTR 6 h2 3bp insert, and miniTTR 6 h3 3bp insert, we generated their DNA templates using primer assembly (42). For each template, primers were generated using the https://primerize.stanford.edu/. The primers were then ordered from Integrated DNA Technologies and solvated in a 100 μM solution of 1x IDTE pH 8.0 buffer. The miniTTR TLR mutants were ordered as four separate oPools from IDT. Upon arrival, they were solvated in 50 μL of 1x IDTE pH 8.0 buffer.

### Generating DNA single templates from primer assembly

The sequence of each primer is listed in the Supplemental Document “Primers.xlsx.” Additionally, the primers for each DNA template are summarized under the primer assembly tab. First, the middle primers (P2-P5 for miniTTR 6 wildtype and P2-P3 for all other templates) were combined and diluted to 1 μM concentration in 100 μL solution. Then, the PCR reaction was mixed by combining 25 μL of Platinum II Hot-Start PCR master mix (ThermoFisher #14000012), two μL of the diluted middle primer solution, two μL of the corresponding forward and reverse primers (P1 and P6 for miniTTR 6 wildtype and P1 and P4 for all other templates) at 100 μM concentration, and 19 μL of RNase-free UltraPure water (ThermoFisher #10977015). The reaction was run in the thermocycler with an initial denaturation for 2 minutes at 94 °C, followed by 30 cycles of denaturation, annealing, and extension steps done for 30 seconds each at 94 °C, 60 °C, and 72°C, respectively. Lastly, a final extension of 5 minutes at 72 °C was performed. The PCR products were then run on a 2% agarose gel for 1 hour at 150 V, and bands of the correct size were extracted and purified using a Zymoclean Gel DNA Recovery Kit (Genesee Scientific #11-301C).

### Generating pooled DNA templates from oligonucleotide pools

The four miniTTR TLR mutant libraries were generated from single-stranded oligonucleotide “Opools” from Integrated DNA Technologies (IDT). The oligo pools are delivered dry and solvated using 50 μL of IDTE pH 8.0 (IDT #11-05-01-13). All libraries were amplified using the same forward and reverse primers (forward: TTCTAATACGACTCACTATAGG, reverse: GTTGTTGTTGTTGTTTCTTT). The primers were also ordered from IDT but were delivered at 100 μM concentration in IDTE pH 8.0 solution. Before setting up the PCR reaction, primers were diluted to 10 μM concentration. The PCR reaction was mixed by combining 25 μL of Q5 High-Fidelity DNA Polymerase (New England Biolabs #M0494S), 2 μL of the oligo pool solution, 2.5 μL of the diluted forward and reverse primers, and 18 μL of RNase-free UltraPure water (ThermoFisher #10977015). The reaction was run in a thermocycler for 20 cycles. Denaturation, annealing, and extension were done at 98 °C, 62 °C, and 72 °C, respectively. The denaturation step was done for 10 seconds, while annealing and extension were done for 15 seconds. An initial denaturation for 30 seconds at 98 °C and a final extension for 5 minutes at 72 °C were also done. The PCR products were then run on a 2% agarose gel for 1 hour at 150 V, and bands of the correct size were extracted and purified using a Zymoclean Gel DNA Recovery Kit (Genesee Scientific #11-301C).

### *In vitro* transcription of RNA constructs

All constructs were transcribed in vitro from DNA into RNA. Before setting up the reaction, the following reagents were prepared: 10x Transcription (Tx) Buffer (400 mM tris-base, 10 mM spermidine, 0.1% Triton X), 25 mM NTP solution, 50 mM DTT, and 250 mM MgCl_2_. Transcription reaction was prepared by combining 10 μL of 10x Tx Buffer, 5 μL of 50mM DTT, 16 μL of 25 mM NTP solution, 8 μL of 250 mM MgCl_2_, 4 μL of t7 polymerase (New England Biolabs #M0251S), 33 μL RNase-Free water, and 24 μL of purified DNA. The concentration of purified DNA was measured via nanodrop and adjusted to 0.3 μM before being added to the transcription reaction. Finally, the reaction mixture was incubated in the thermocycler at 37 °C for 6 hours. After the reaction, the DNA template was digested using DNase I, and it was included in the RNA Clean and Concentrator-5 with the DNase I kit (Genesee Scientific #R1014). The RNA was then purified using the same kit. Before proceeding to the DMS MaPseq experiment, the concentration of RNA was measured through its absorbance value on the nanodrop. Additionally, we verified RNA length by running the sample through a 4% agarose denaturing gel for 1 hour at 150 volts.

### DMS-MaPseq protocol

DMS-MaPseq was done on all purified RNA constructs. The specific conditions for RNA folding and RNA modification with DMS varied accordingly for the various buffer titrations and magnesium titrations that were done. First, we measured 10 pmol of RNA in 5 μL of RNase-Free water for each reaction condition. RNA was denatured in the thermocycler at 90 °C for 4 minutes and immediately cooled for 3 minutes at 4 °C. Then the RNA was pipetted into a folding solution containing 16.5 μL of buffer (at reach the desired concentration) and 1 μL of MgCl_2_ at the desired concentrations. For example, for the condition specifying 50 mM sodium cacodylate and 10 mM Mg^2+^, 16.5 μL of 76 mM sodium cacodylate was mixed with 1 μL of 250 mM MgCl_2_ to prepare the reach a final concentration of 50 mM sodium cacodylate and 10 mM MgCl_2_. All conditions were done similarly to prepare a final reaction volume of 25 uL. The RNA was allowed to fold in solution for 30 minutes at room temperature. During this time, a DMS solution was prepared in the fume hood. This was done by adding 15 μL of DMS (Sigma-Aldrich #D186309) to 85 μL of 100% ethanol (Decon Labs cat. #2716). Once RNA was folded for 30 minutes, 2.5 μL of DMS solution was added to the RNA buffer solution to modify the RNA. The reaction was quenched 6 minutes after DMS with 25 μL of BME (ThermoFisher cat. #125470010) to the solution. Then, the modified RNA was purified using the RNA Clean & Concentrator-5 kit (Genesee Scientific #R1014). The RNA sample was eluted from the column with 7 μL of RNase-Free water. The concentration of RNA was measured using the Qubit RNA BR Assay Kit (ThermoFisher #Q10211). 1 μL of RNA sample was used to measure the concentration of modified RNA.

To read out the methylations of A and C, we utilized TGIRT III reverse transcriptase, which incorporates mutations of the cDNA strand during reverse transcription. Before setting up the reverse transcription reaction, the following solutions were prepared: 5x TGIRT buffer (250 mM Tris-HCl pH 8.3, 375 mM KCl, 15 mM MgCl2), 10 mM dNTPs, 100 mM DTT, 0.4 M NaOH, and Quench Acid (1.43 M NaCl, 0.57 M HCl, 1.29 M sodium acetate). Additionally, the modified RNA sample was diluted to a concentration of 0.25 μM and one RTB primer to introduce a barcode for demultiplexing (**Supplemental Document Primers.xslx**) to 0.285 μM. Finally, the reverse transcription reaction was mixed by combining 2.4 μL of 5x TGIRT buffer, 1.2 μL of 10 mM dNTP solution, 0.6 μL of 100 mM DTT, 0.5 μL of TIGIRT-III enzyme, 6.4 μL of 0.25 μM modified RNA, and 1 μL of 0.285 μM RTB primer. The reaction was mixed thoroughly and then incubated at 57C for 2 hours. After incubation, 5 μL of 0.4 M NaOH was added to the reaction products. The sample was heated in the thermocycler at 90 °C for 4 minutes and then cooled immediately for 3 minutes at 4C. Then, 2.5 μL of quench acid was added to neutralize the NaOH. Note that the exact volume of quench acid required to neutralize the NaOH will vary slightly for different batches. Next, the cDNA product was purified using the Oligo Clean and Concentrator Kit (Genesee Scientific #11-380B). Before starting the purification, 30 μL of RNase-Free water was added to the reaction products to bring the volume to 50 μL. The cDNA was eluted from the column using 15 μL of RNase-free water.

Then, the cDNA was amplified with PCR, similar to how the original DNA templates were amplified from an oligo pool. The PCR used forward and reverse primers ordered from IDT dissolved in IDTE pH 8 buffer at a concentration of 100 μM. All constructs were amplified with the same FWD primer AATGATACGGCGACCACCGAGATCTACACTCTTTCCCTACACGCGCTCTTCCG. The minittr-6 was amplified with reverse primer CAAGCAGAAGACGGCATACGAGATCGGTCTCGGCATTCCTGCTGAACCGCTCTTCCGATCTGGTACTATGTACCAAAG while all others were amplified with reverse primer CAAGCAGAAGACGGCATACGAGATCGGTCTCGGCATTCCTGCTGAACCGCTCTTCCGATCTGGAACAGCACTTCGGTGCAAA. These primers were diluted in a 1:10 dilution to a concentration of 10 uM before mixing the reaction. To mix the reaction, 25 μL of Q5 High-Fidelity DNA Polymerase (New England Biolabs #M0494S), 2.5 μL of diluted forward and reverse primer, 2.0 μL of purified cDNA, and 18 μL of RNase-Free water were combined. The reaction was run in a thermocycler for 16 cycles. Denaturation, annealing, and extension were done at 98C, 62C, and 72 °C, respectively. The denaturation step was done for 10 seconds, while annealing and extension were done for 15 seconds. An initial denaturation for 30 seconds at 98C and a final extension for 5 minutes at 72 °C were also done. PCR products were gel purified using the E-Gel Power Snap Plus Electrophoresis System (ThermoFisher #G9301). 5 μL of PCR products was run on a 2% E-gel EX Agarose Gel (ThermoFisher #G401002) for 10 minutes according to the settings programmed into the gel system. Then, the gel was imaged, and bands of the correct size were excised carefully using a metal spatula. The DNA was then purified using the Zymoclean Gel DNA Recovery Kit (Genesee Scientific #11-301C). Lastly, the purified DNA concentration was measured using the Qubit 1X dsDNA High Sensitivity Assay Kit (ThermoFisher #Q33230).

### DMS-MaPseq data analysis

Sequencing was performed on either an Iseq 100 or Novaseq 6000. Each sequencing run was first demultiplexed using the RTB barcodes added during the RT. Demultiplexing was performed using novobarcode (https://www.novocraft.com/documentation/novobarcode/demultiplexing-barcodedindexed-reads-with-novobarcode/).

`novobarcode -b rtb_barcodes.fa -f test_R1_001.fastq test_R2_001.fastq`
where rtb_barcodes.fa gives a list of sequencing barcodes, such as

`Distance 4`

`Format 5`

`RTB021 CCAATGGGTGTA`

`RTB022 AGCCAAAACTGG`

`RTB023 GTGTGTTTGCCC`

Where three barcodes are specified, distance is the distance in base pairs between a barcode and an allowable read. Format specifies that the barcode will be on the 5′ end of the read 1. For the pools that contained TLR mutants, we performed an additional second demultiplexing on the helix under the TLR. These seven base pair helices are unique to each construct in the pool. This second step ensured there were no missed alignments, as even a few percent missed alignments of reads could add additional noise. Demultiplexing of internal helices was performed by custom software developed in-house (https://github.com/jyesselm/barcode_demultiplex).

`barcode_demultiplex -csv constructs.csv -fq1 test_R2_001.fastq -fq2 test_R1_001.fastq -helix 1 0 7 -m 1`

constructs.csv is a comma-delimited formatted file containing the name, sequence, and structure in dot-bracket format of each RNA in a pool. The -fq1 and -fq2 flags take FASTQ formatted sequencing output files, with fastq1 as the file that contains the RNA sequence from 5′ to 3′. -helix specifies which helix to use in occurrence from the 5’ end. -helix 1 0 7 specifies using the second helix (note this is zero-indexed) and uses base pairs 0 to 7. Lastly, -m 1 indicates accepting sequences that are at most hamming distance of 1 away from the target barcode sequence. This command uses SeqKit (43) to find the occurrence of a given barcode in the supplied FASTQ files in the approximate position it should exist. It will ensure the barcode sequence exists in both the forward and reverse reads if possible. barcode_demultiplex outputs reads into separated paired FASTQ formatted files containing only reads with a given barcode identified.

Lastly, we used the RNA mutational profiling (RNA-MaP) tool to count mutations to determine the mutational fraction at each nucleotide position (https://github.com/YesselmanLab/rna_map). This is an open-source tool developed to simplify mutational profiling analysis.

`rna-map -fa test.fasta -fq1 test_mate1.fastq -fq2 test_mate2.fastq -- dot-bracket test.csv --param-preset barcoded-libraries`

The rna-map command requires a FASTA-formatted file containing all DNA reference sequences and the paired sequencing reads generated from the previous step. We also supplied a CSV file containing each RNA’s dot bracket structure with the --dot-bracket flag and applied stricter constraints to how well each sequence needs to align to a read --param-preset barcoded-libraries. All the data are stored in JSON format in a pandas DataFrame.

### Computing [Mg^2+^]½ from 16-point [Mg^2+^] titration

To compute the [Mg^2+^]_½_ of each construct, we took the 16-point [Mg^2+^] titration data and extracted the DMS reactivity for the three adenines in the GAAA tetraloop (*average GAAA DMS*). The first 25 construct library 5 mM [Mg^2+^] points had unexpectedly high reactivity, so they were removed. To estimate [Mg^2+^]_½,_ we computed the relative protection values of each Mg^2+^ concentration *i, which* were calculated below.

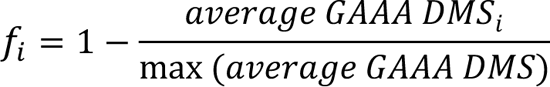

Where *f*_i_ is the relative DMS protection that a given Mg^2+^ concentration imparts compared to the 0 Mg^2+^ concentration condition, this number ranges from 0 (no protection) to 1 (completely protected). We fit these values using a modified Hill equation as used in a previous study (6).

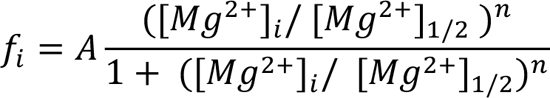

Where [Mg^2+^]_i_ is the Mg^2+^ concentration at a given measurement, A is the scaling factor, and n is the Hill coefficient. These fits were performed with SciPy’s curve_fit function for each data set. One hundred bootstrap steps were performed to estimate the error in the fit.

To compute the Δ*G* of a given tertiary contact formation can be calculated using the following equation.

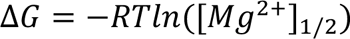

R is the gas constant, and T is the temperature at 20 °C.

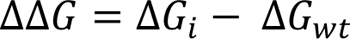

Computing ΔΔ*G* can be done by subtracting any computed Δ*G* from the Δ*G*_’(_, which is the Δ*G* of the wild type.

### Performing solvent accessibility calculations

We computed solvent accessibility of the N1 position of A4, A5, and A8 over the three structures representing our three states. We used PDBs 1TLR, 1GID (removed GAAA tetraloop), and 1GID to represent our three states. We used freesasa (https://freesasa.github.io/python/), using the Lee Richards model with a probe radius of 1.0 Å.

### Generating 3D model for TLR mutants with FARFAR2

Hypothesizing that their binding orientation would closely resemble that of the wild type, given their similar thermodynamic profiles. In all models, we used the orientation from the wild-type to constrain the binding mode for each TLR mutation. This was performed by supplying the original 1C-12G pair and the G-C pair that closes the GAAA tetraloop from the crystal structure of the P4-P6 domain of the tetrahymena ribozyme (PDB: 1GID). This was performed using the `rna_denovo` command in Rosetta, which runs FARFAR2.

`rna_denovo -fasta test.fasta -secstruct_file test.secstruct_file -s start.pdb -minimize_rna true -nstruct 1000`

Where the `-fasta` specifies a FASTA formatted file containing the sequences of the three strands. Like the example below.

`>loop ggaaac`

`>receptor_1 gauaugg`

`>receptor_2 ccuaaaguc`

`-secstruct_file` specifies a secondary structure file that contains the dot bracket of the secondary structure matched with the primary sequence found in the FASTA file. An example is shown below.

`(….)+(((..((+))….)))`

ggaaac+gauaugg+ccuaaaguc

`-s` supplies the coordinate file containing the 1C-12G pair and the G-C pair that closes the GAAA tetraloop. `-minimize_rna true` flag does a round steepest decent minimization after generating each model. Lastly, `-nstruct 1000`, generates 1000 individual models. All models are stored in default.out silent formatted file. To select the top 10 lowest energy models, we used the `extract_lowscore_decoys.py` script part of Rosetta tools (https://www.rosettacommons.org/docs/latest/application_documentation/tools/Tools).

`extract_lowscore_decoys.py default.out 10`

## Notes

http://doi.org/10.6084/m9.figshare.25331758

https://github.com/jyesselm/q_dms_ttr_paper

https://www.ncbi.nlm.nih.gov/bioproject/PRJNA1086549

### Competing Interest Statement

The authors have declared no competing interest.

## References

1. Ban N, Nissen P, Hansen J, Moore PB, Steitz TA. The complete atomic structure of the large ribosomal subunit at 2.4 Å resolution. Science. 2000;289(5481):905–20.

2. Schmitzová J, Cretu C, Dienemann C, Urlaub H, Pena V. Structural basis of catalytic activation in human splicing. Nature. 2023;617(7962):842-+.

3. Han JH, Shyamala V, Richman KH, Brauer MJ, Irvine B, Urdea MS, et al. Characterization of the Terminal Regions of Hepatitis-C Viral-Rna - Identification of Conserved Sequences in the 5’ Untranslated Region and Poly(a) Tails at the 3’ End. P Natl Acad Sci USA. 1991;88(5):1711–5.

4. Lu K, Heng X, Summers MF. Structural Determinants and Mechanism of HIV-1 Genome Packaging. J Mol Biol. 2011;410(4):609–33.

5. Jasinski D, Haque F, Binzel DW, Guo P. Advancement of the Emerging Field of RNA Nanotechnology. ACS Nano. 2017;11(2):1142–64.

6. Yesselman JD, Eiler D, Carlson ED, Gotrik MR, d’Aquino AE, Ooms AN, et al. Computational design of three-dimensional RNA structure and function. Nat Nanotechnol. 2019;14(9):866-+.

7. Nissen P, Ippolito JA, Ban N, Moore PB, Steitz TA. RNA tertiary interactions in the large ribosomal subunit: The A-minor motif. P Natl Acad Sci USA. 2001;98(9):4899–903.

8. Butcher SE, Pyle AM. The Molecular Interactions That Stabilize RNA Tertiary Structure: RNA Motifs, Patterns, and Networks. Accounts Chem Res. 2011;44(12):1302–11.

9. Young BT, Silverman SK. The GAAA tetraloop-receptor interaction contributes differentially to folding thermodynamics and kinetics for the P4-P6 RNA domain. Biochemistry-Us. 2002;41(41):12271–6.

10. Bokinsky G, Rueda D, Misra VK, Rhodes MM, Gordus A, Babcock HP, et al. Single-molecule transition-state analysis of RNA folding. Proc Natl Acad Sci U S A. 2003;100(16):9302–7.

11. Rangan P, Masquida B, Westhof E, Woodson SA. Assembly of core helices and rapid tertiary folding of a small bacterial group I ribozyme. Proc Natl Acad Sci U S A. 2003;100(4):1574–9.

12. Hodak JH, Downey CD, Fiore JL, Pardi A, Nesbitt DJ. Docking kinetics and equilibrium of a GAAA tetraloop-receptor motif probed by single-molecule FRET. Proc Natl Acad Sci U S A. 2005;102(30):10505–10.

13. Chauhan S, Woodson SA. Tertiary interactions determine the accuracy of RNA folding. J Am Chem Soc. 2008;130(4):1296–303.

14. Draper DE. RNA folding: thermodynamic and molecular descriptions of the roles of ions. Biophys J. 2008;95(12):5489–95.

15. Sattin BD, Zhao W, Travers K, Chu S, Herschlag D. Direct measurement of tertiary contact cooperativity in RNA folding. J Am Chem Soc. 2008;130(19):6085–7.

16. Holmstrom ED, Fiore JL, Nesbitt DJ. Thermodynamic origins of monovalent facilitated RNA folding. Biochemistry-Us. 2012;51(18):3732–43.

17. Bisaria N, Greenfeld M, Limouse C, Mabuchi H, Herschlag D. Quantitative tests of a reconstitution model for RNA folding thermodynamics and kinetics. Proc Natl Acad Sci U S A. 2017;114(37):E7688–E96.

18. White NA, Hoogstraten CG. Thermodynamics and kinetics of RNA tertiary structure formation in the junctionless hairpin ribozyme. Biophys Chem. 2017;228:62–8.

19. Bonilla SL, Denny SK, Shin JH, Alvarez-Buylla A, Greenleaf WJ, Herschlag D. High-throughput dissection of the thermodynamic and conformational properties of a ubiquitous class of RNA tertiary contact motifs. P Natl Acad Sci USA. 2021;118(33).

20. Denny SK, Bisaria N, Yesselman JD, Das R, Herschlag D, Greenleaf WJ. High-Throughput Investigation of Diverse Junction Elements in RNA Tertiary Folding. Cell. 2018;174(2):377-+.

21. Shin JH, Bonilla SL, Denny SK, Greenleaf WJ, Herschlag D. Dissecting the energetic architecture within an RNA tertiary structural motif via high-throughput thermodynamic measurements. P Natl Acad Sci USA. 2023;120(11).

22. Cordero P, Kladwang W, VanLang CC, Das R. Quantitative Dimethyl Sulfate Mapping for Automated RNA Secondary Structure Inference. Biochemistry-Us. 2012;51(36):7037–9.

23. Deigan KE, Li TW, Mathews DH, Weeks KM. Accurate SHAPE-directed RNA structure determination. P Natl Acad Sci USA. 2009;106(1):97–102.

24. Tomezsko P, Swaminathan H, Rouskin S. Viral RNA structure analysis using DMS-MaPseq. Methods. 2020;183:68–75.

25. Tomezsko P, Swaminathan H, Rouskin S. DMS-MaPseq for Genome-Wide or Targeted RNA Structure Probing In Vitro and In Vivo. Methods Mol Biol. 2021;2254:219–38.

26. Zubradt M, Gupta P, Persad S, Lambowitz AM, Weissman JS, Rouskin S. DMS-MaPseq for genome-wide or targeted RNA structure probing. Nat Methods. 2017;14(1):75–82.

27. Haddrick M, Lear AL, Cann AJ, Heaphy S. Evidence that a kissing loop structure facilitates genomic RNA dimerisation in HIV-1. J Mol Biol. 1996;259(1):58–68.

28. Paillart JC, Westhof E, Ehresmann C, Ehresmann B, Marquet R. Non-canonical interactions in a kissing loop complex: the dimerization initiation site of HIV-1 genomic RNA. J Mol Biol. 1997;270(1):36–49.

29. Tijerina P, Mohr S, Russell R. DMS footprinting of structured RNAs and RNA-protein complexes. Nat Protoc. 2007;2(10):2608–23.

30. Lan TCT, Allan MF, Malsick LE, Woo JZ, Zhu C, Zhang F, et al. Secondary structural ensembles of the SARS-CoV-2 RNA genome in infected cells. Nat Commun. 2022;13(1):1128.

31. Fiore JL, Hodak JH, Piestert O, Downey CD, Nesbitt DJ. Monovalent and divalent promoted GAAA tetraloop-receptor tertiary interactions from freely diffusing single-molecule studies. Biophys J. 2008;95(8):3892–905.

32. Takamoto K, Das R, He Q, Doniach S, Brenowitz M, Herschlag D, et al. Principles of RNA compaction: insights from the equilibrium folding pathway of the P4-P6 RNA domain in monovalent cations. J Mol Biol. 2004;343(5):1195–206.

33. Butcher SE, Dieckmann T, Feigon J. Solution structure of a GAAA tetraloop receptor RNA. Embo J. 1997;16(24):7490–9.

34. Cate JH, Hanna RL, Doudna JA. A magnesium ion core at the heart of a ribozyme domain. Nat Struct Biol. 1997;4(7):553–8.

35. Basu S, Rambo RP, Strauss-Soukup J, Cate JH, Ferré-D’Amaré AR, Strobel SA, et al. A specific monovalent metal ion integral to the AA platform of the RNA tetraloop receptor. Nat Struct Biol. 1998;5(11):986–92.

36. Das R, Karanicolas J, Baker D. Atomic accuracy in predicting and designing noncanonical RNA structure. Nat Methods. 2010;7(4):291–4.

37. Watkins AM, Rangan R, Das R. FARFAR2: Improved Rosetta Prediction of Complex Global RNA Folds. Structure. 2020;28(8):963-+.

38. Yesselman JD, Das R. Modeling Small Noncanonical RNA Motifs with the Rosetta FARFAR Server. Methods Mol Biol. 2016;1490:187–98.

39. Homan PJ, Favorov OV, Lavender CA, Kursun O, Ge X, Busan S, et al. Single-molecule correlated chemical probing of RNA. Proc Natl Acad Sci U S A. 2014;111(38):13858–63.

40. Cheng CY, Kladwang W, Yesselman JD, Das R. RNA structure inference through chemical mapping after accidental or intentional mutations. Proc Natl Acad Sci U S A. 2017;114(37):9876–81.

41. Mustoe AM, Weidmann CA, Weeks KM. Single-Molecule Correlated Chemical Probing: A Revolution in RNA Structure Analysis. Acc Chem Res. 2023;56(7):763–75.

42. Tian S, Yesselman JD, Cordero P, Das R. Primerize: automated primer assembly for transcribing non-coding RNA domains. Nucleic Acids Research. 2015;43(W1):W522–W6.

43. Shen W, Le S, Li Y, Hu F. SeqKit: A Cross-Platform and Ultrafast Toolkit for FASTA/Q File Manipulation. PLoS One. 2016;11(10):e0163962.

