## Supplemental Document for "High-throughput determination of RNA tertiary contact thermodynamics by quantitative DMS chemical mapping"

### Supplemental material for High-throughput determination of RNA tertiary contact thermodynamics by quantitative DMS chemical mapping

Bret Lange[1], Ricardo G. Gil[1], and Joseph D. Yesselman[1]*

[1]Department of Chemistry, University of Nebraska, 639 North 12$^{th}$ St, Lincoln, NE 68588, USA

### Supplemental Figures

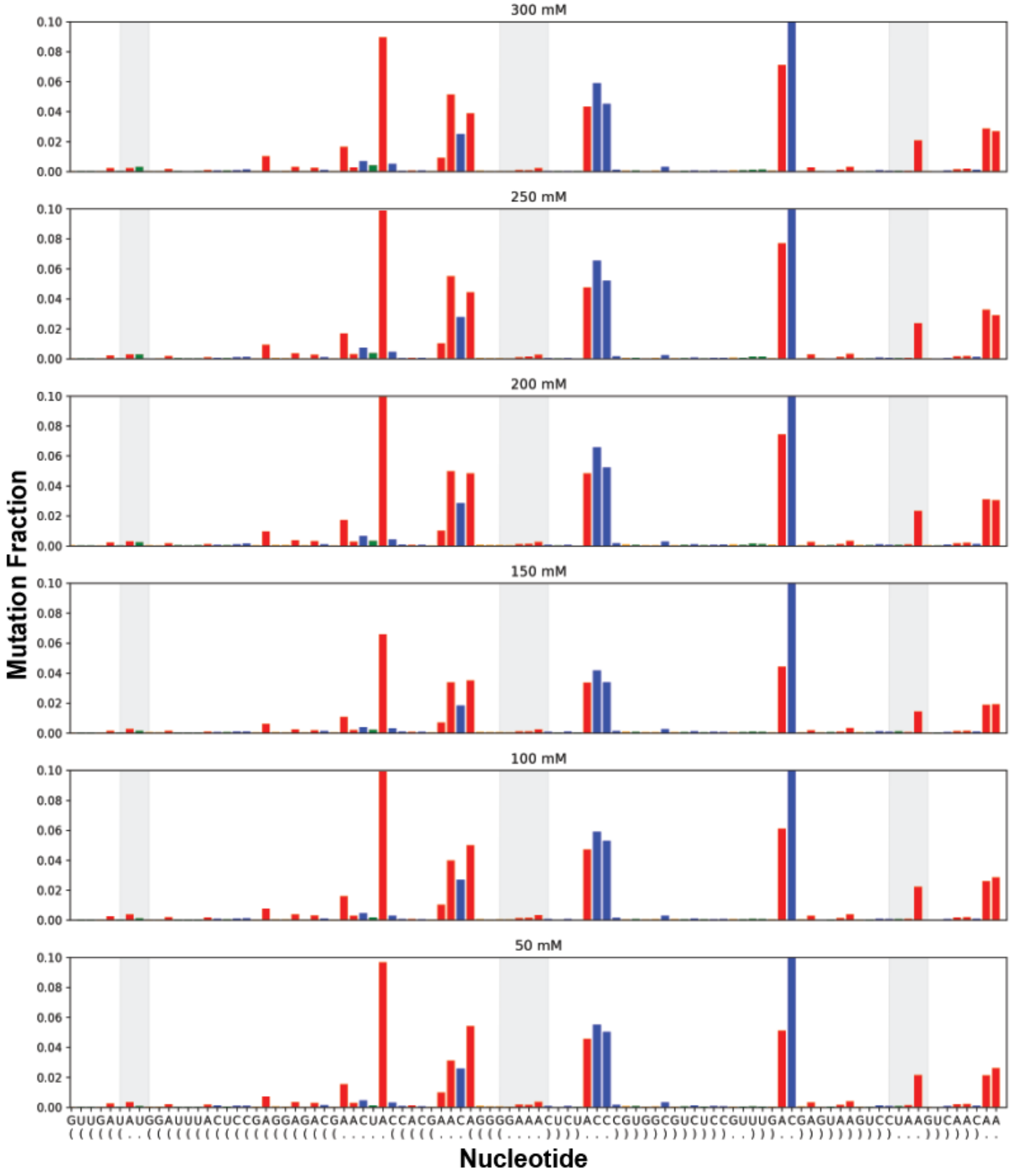

Supplemental Figure 1: DMS-MaPseq mutation fractions as a function of sodium cacodylate buffer titration with 10 mM $Mg^{2+}$.

Gray highlights the tetraloop in the center and the tetraloop receptor on the left and right.

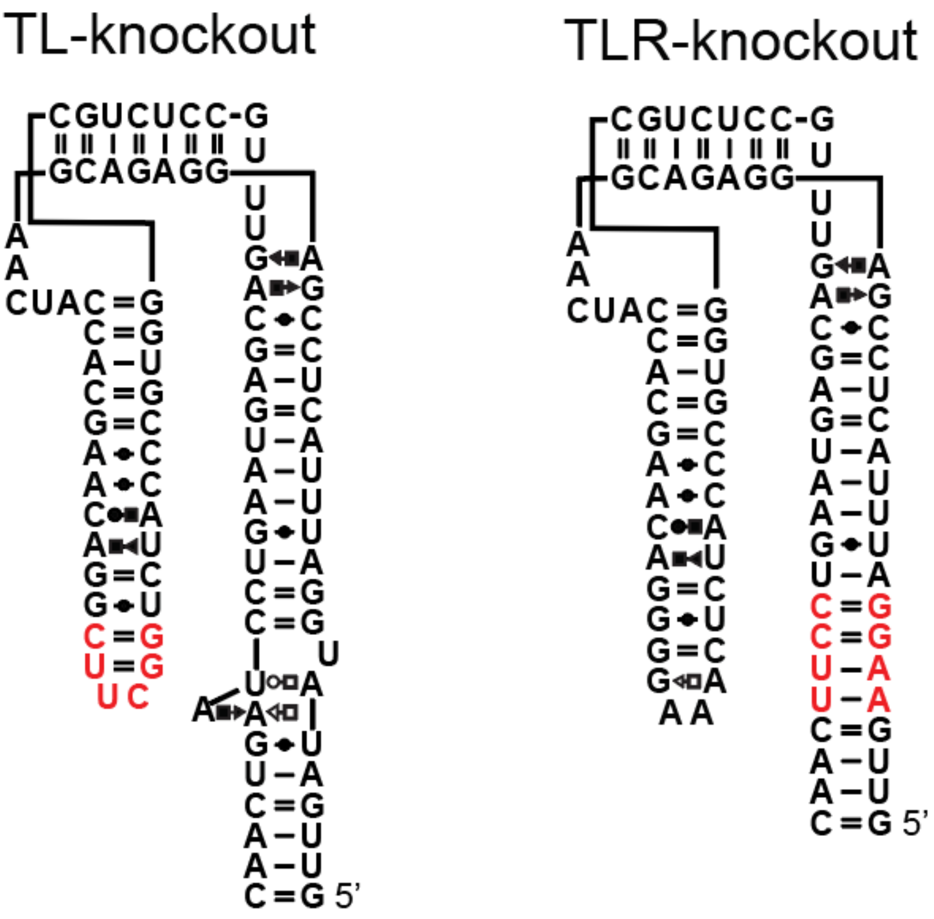

Supplemental Figure 2: Secondary structure of TL-knockout and TLR-knockout

Red denotes mutations. In TL-knockout mutated the GAAA tetraloop to UUCG, in the TLR-knockout mutated the TLR to a Watson-Crick helix.

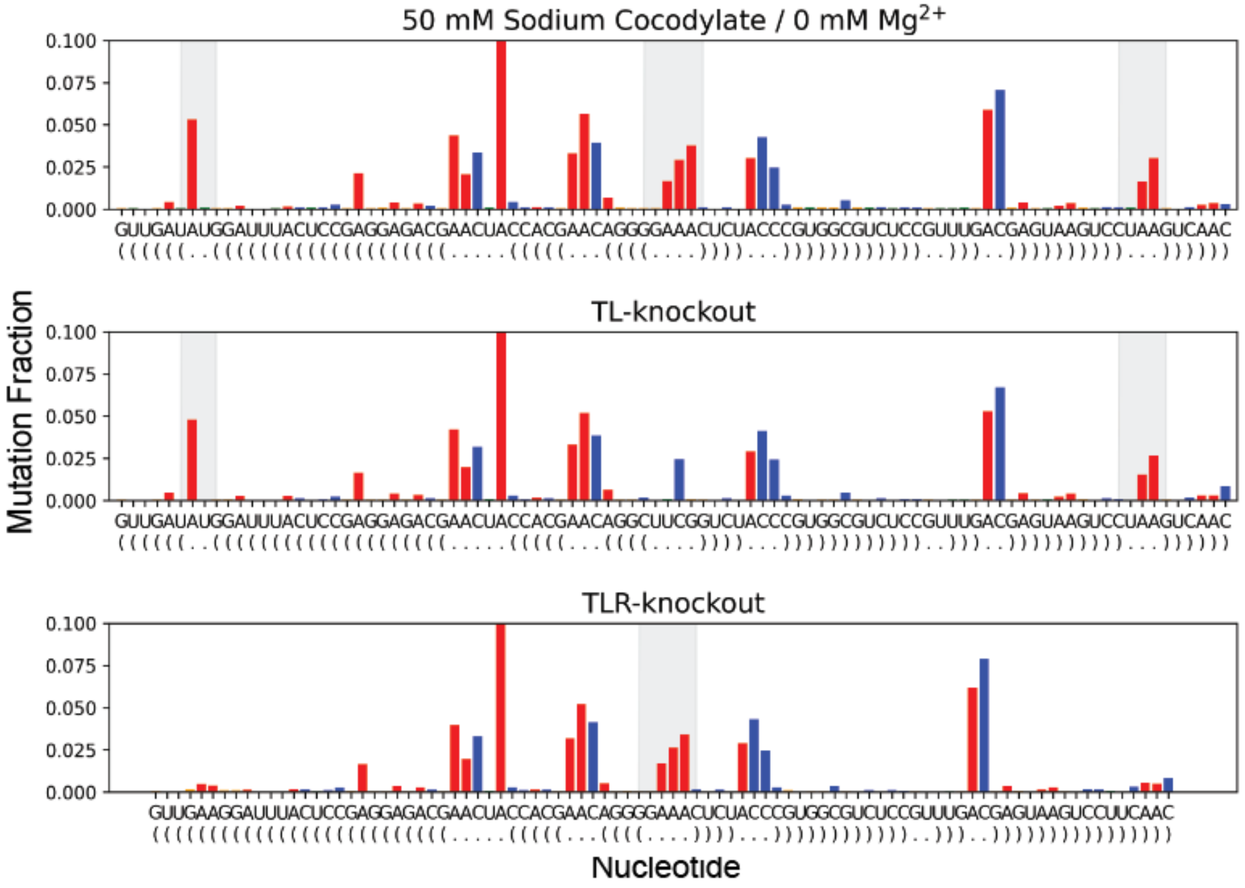

Supplemental Figure 3: 50 mM sodium cacodylate correctly captures the unformed state of the tetraloop/ tetraloop receptor contact.

The wild-type without Mg2+ ions with 50 mM sodium cacodylate. The highlights are the tetraloop and tetraloop-receptor. Their reactivity closely matches the TL-knockout and TLR-knockout constructs.

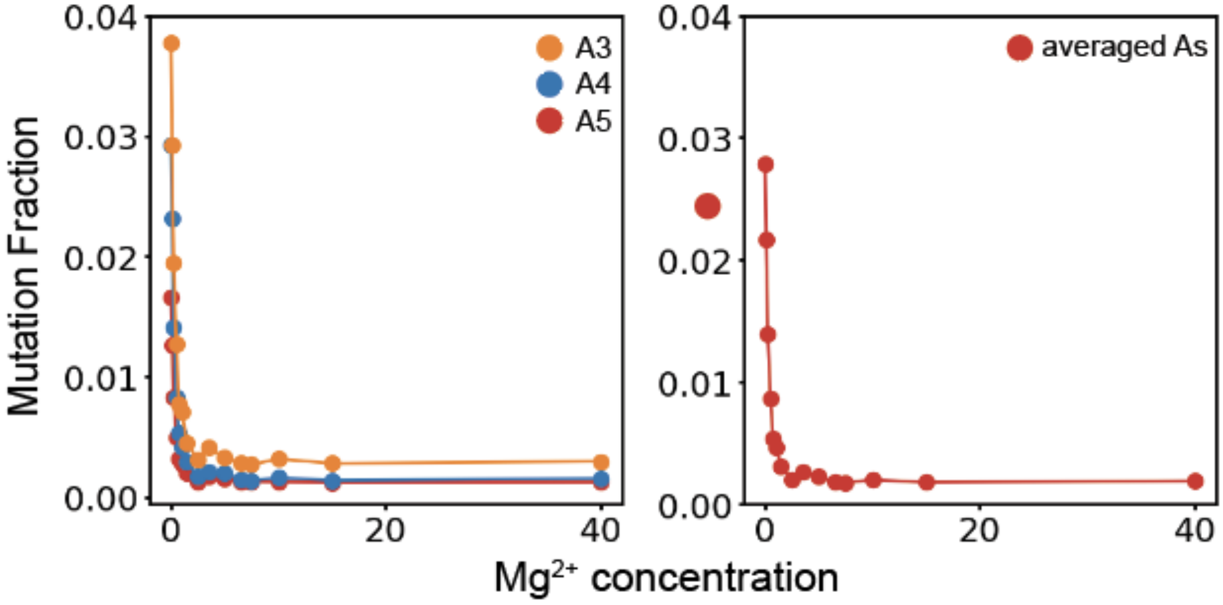

Supplemental Figure 4: The three As in GAAA have similar reactivity profiles and can be averaged.

(Left) Shows the three As in the GAAA loop. Each has a near-identical reactivity profile as a function of $Mg^{2+}$. (Right) Averaging the three As simplifies analysis and fitting.

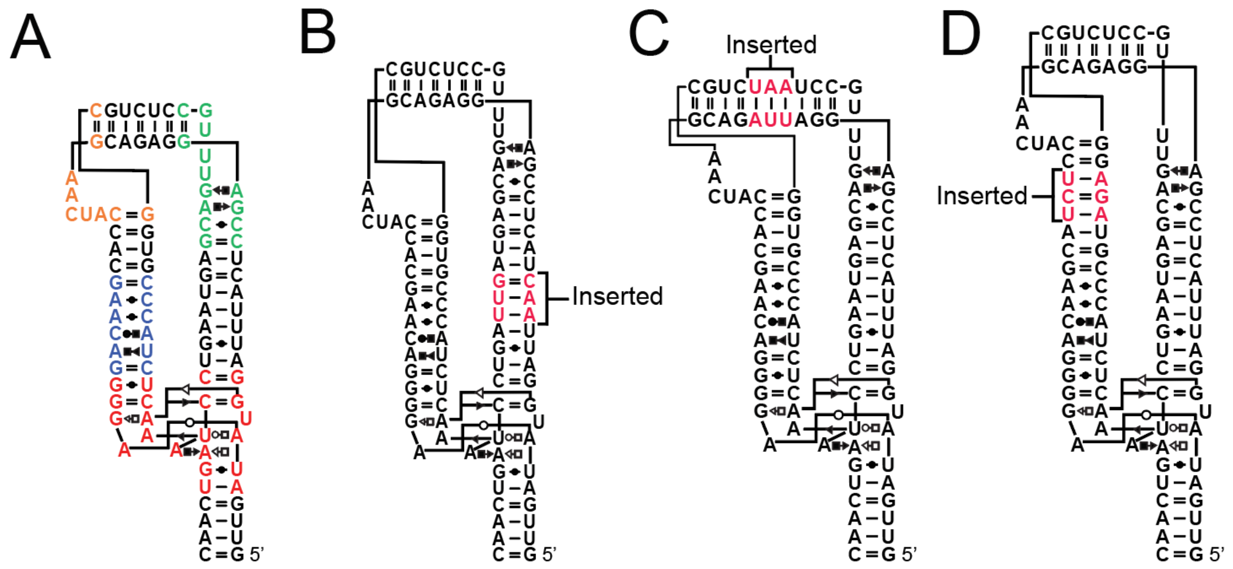

Supplemental Figure 6: The three different destabilizing mutations

Shows the three different 3 bp insertions to destabilize the TL/TLR tertiary contact. (B) Insertion into helix 1 (H1). (C) Insertion into helix 2 (H2). (D) Insertion into helix 3 (H3).

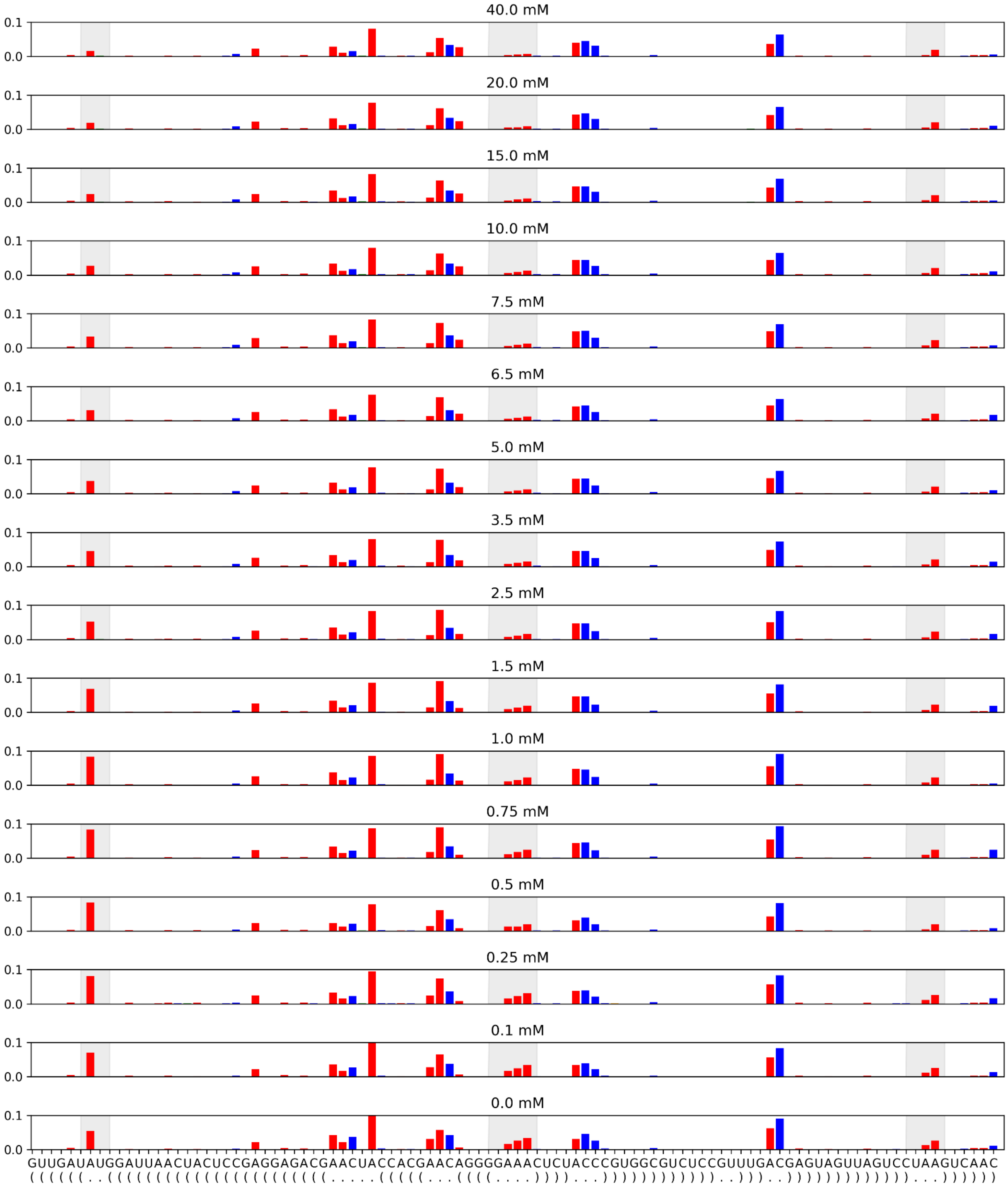

Supplemental Figure 7: DMS-MaPseq mutation fractions for H1 insertion as a function of $Mg^{2+}$ titration.

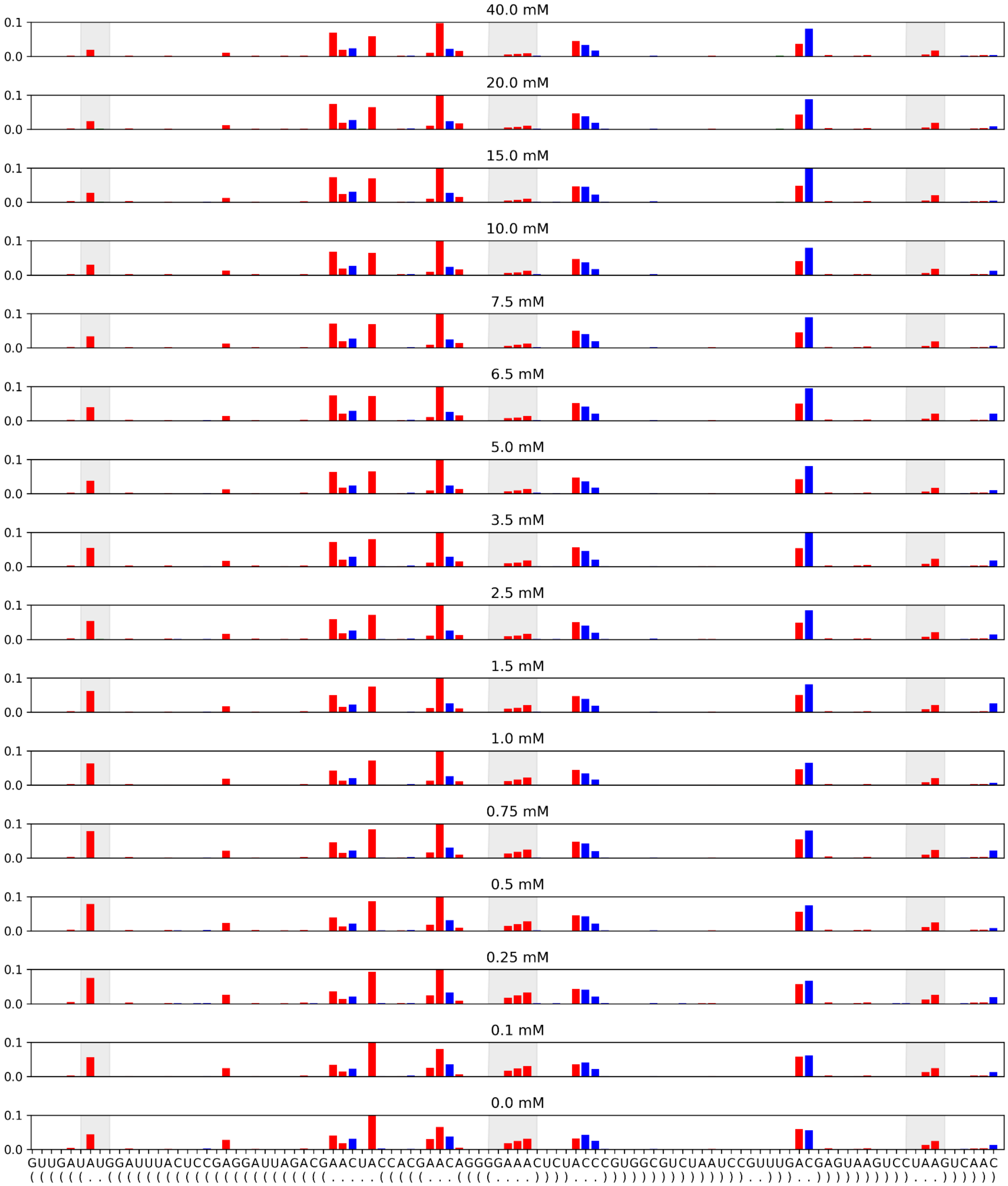

Supplemental Figure 8: DMS-MaPseq mutation fractions for H2 insertion as a function of $Mg^{2+}$ titration.

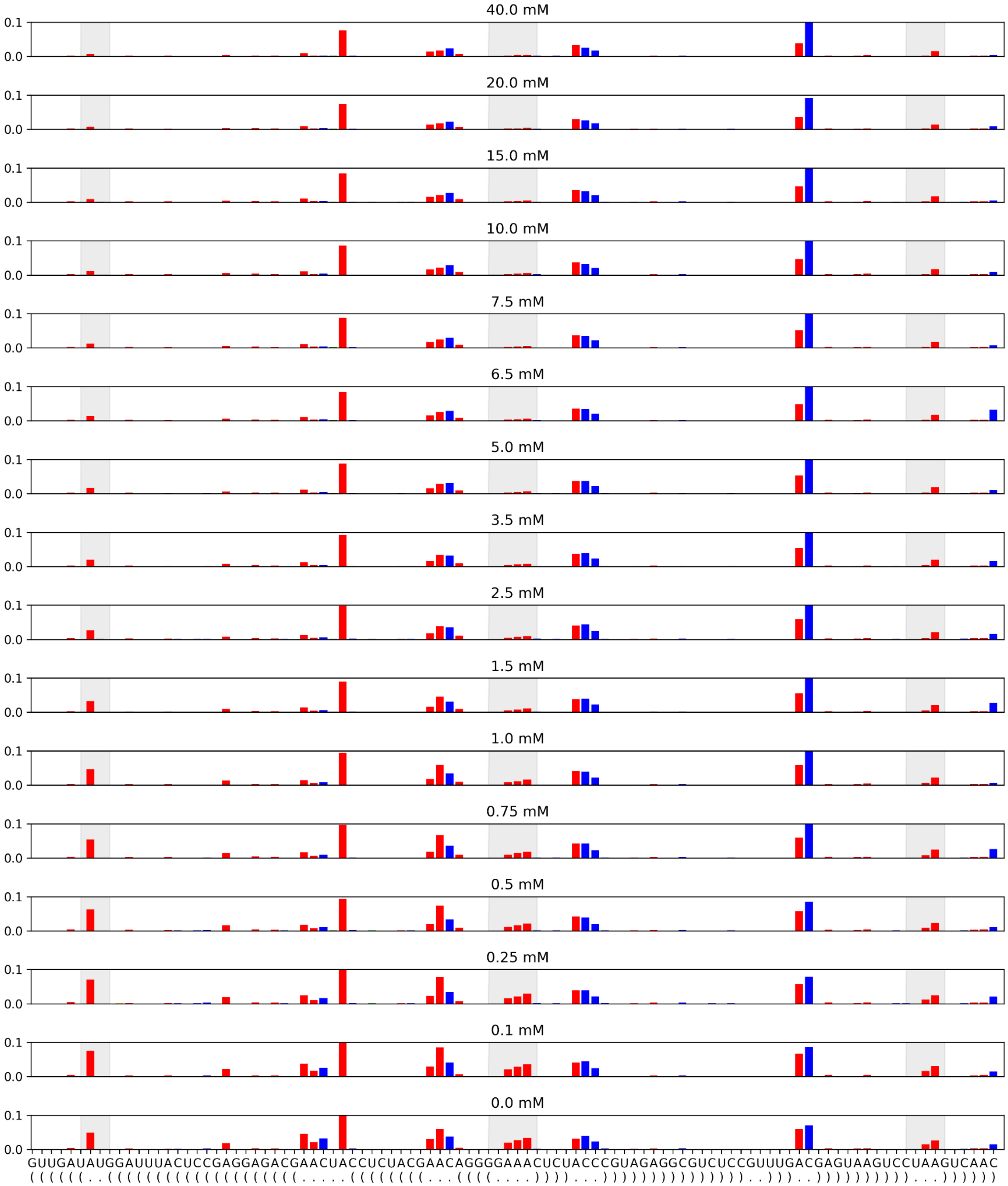

Supplemental Figure 9: DMS-MaPseq mutation fractions for H3 insertion as a function of $Mg^{2+}$ titration.

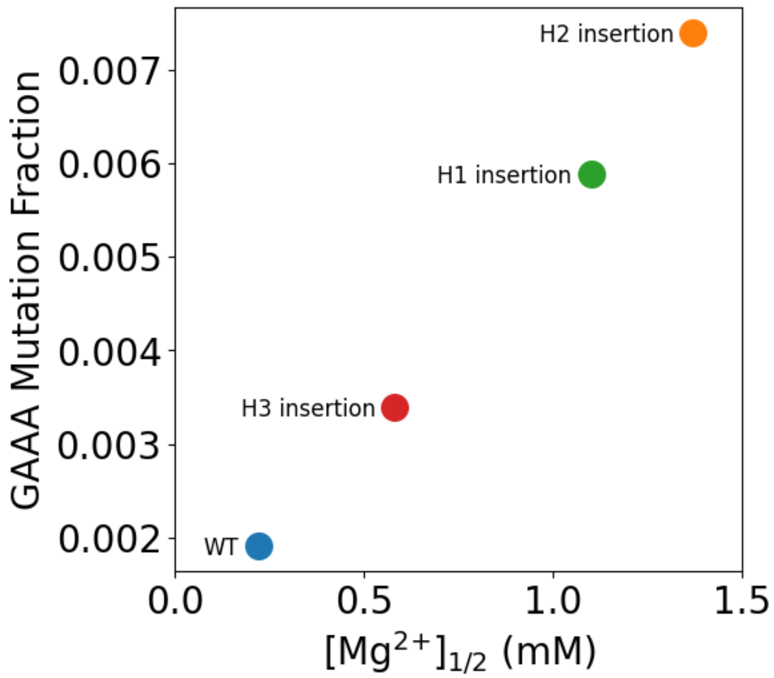

Supplemental Figure 10: Direct correlation between GAAA mutation fraction at 40 mM and $Mg^{2+}$.

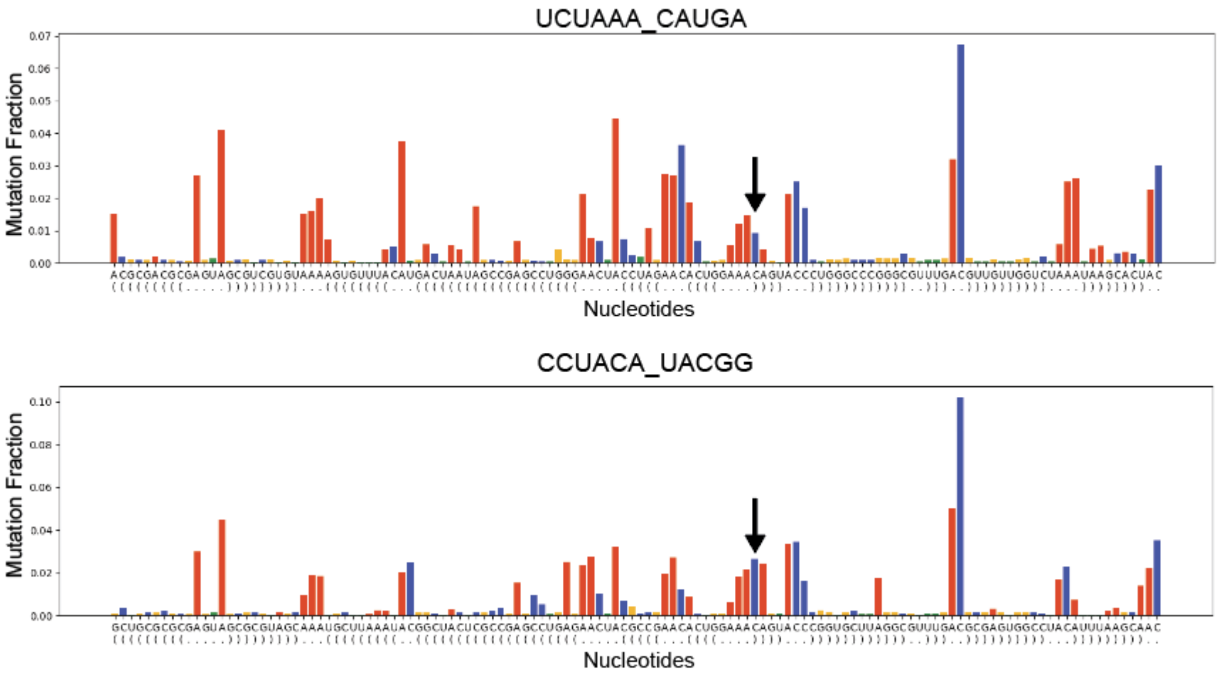

Supplemental Figure 11: GC flanking pair breaking open in UCUAAA_CAUGA and CCUACA_UACGG

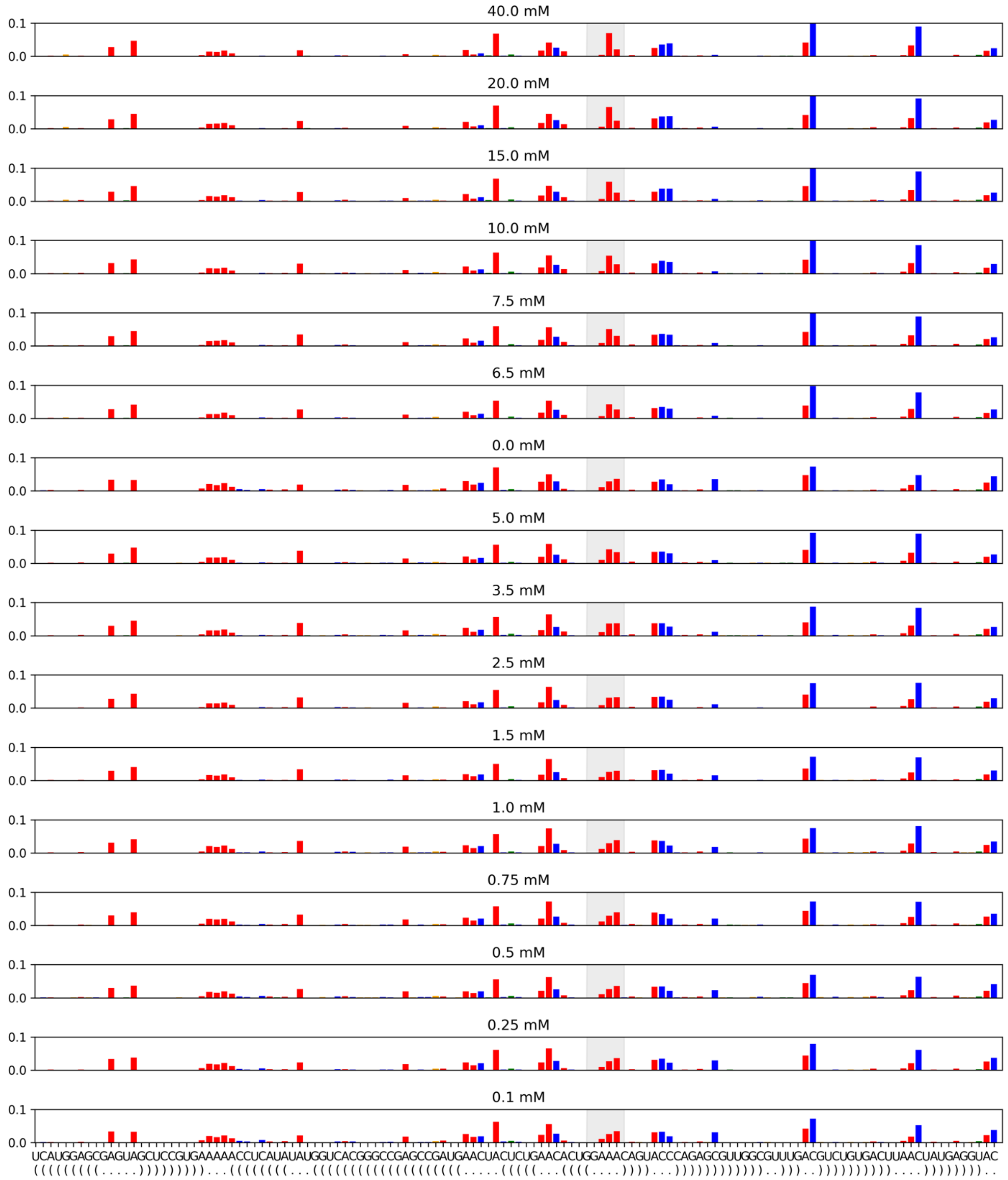

Supplemental Figure 12: CUUAAC_UAUGG titration does not have the GAAA reactivity decrease.

Instead, only A3 and A5 go down while A2 increases in reactivity which is not consistent with our model of TL/TLR binding.

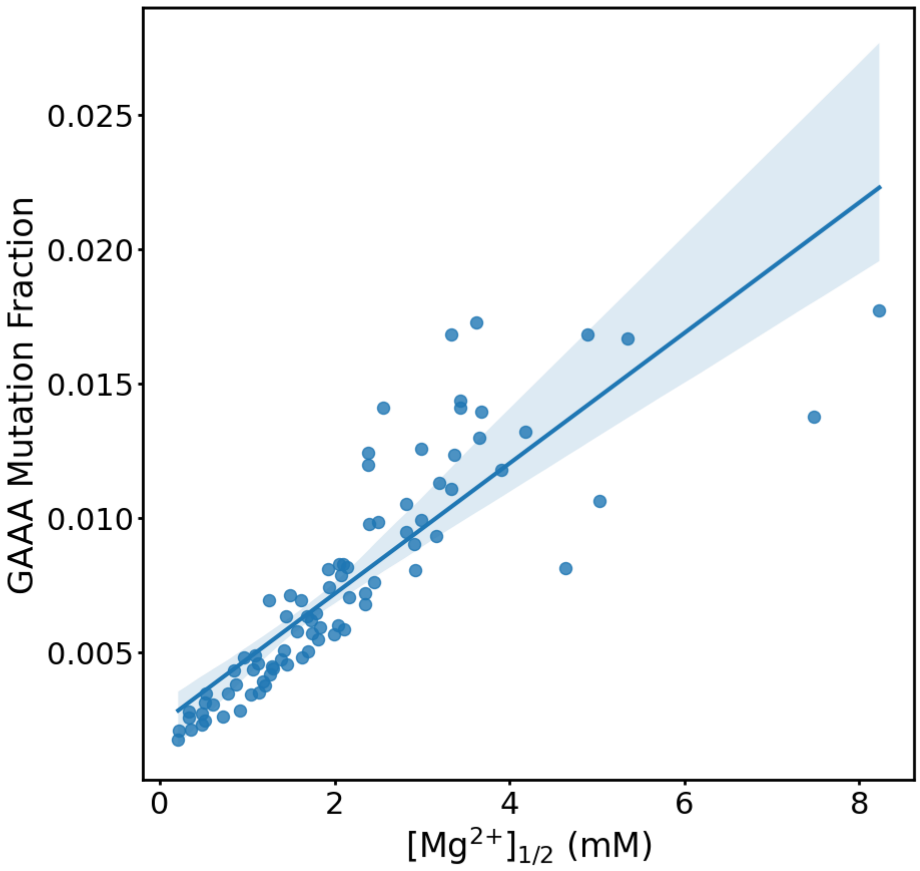

Supplemental Figure 13: Strong correlation between $[Mg^{2+}]_{1/2}$ and GAAA reactivity at 7.5 mM $Mg^{2+}$.

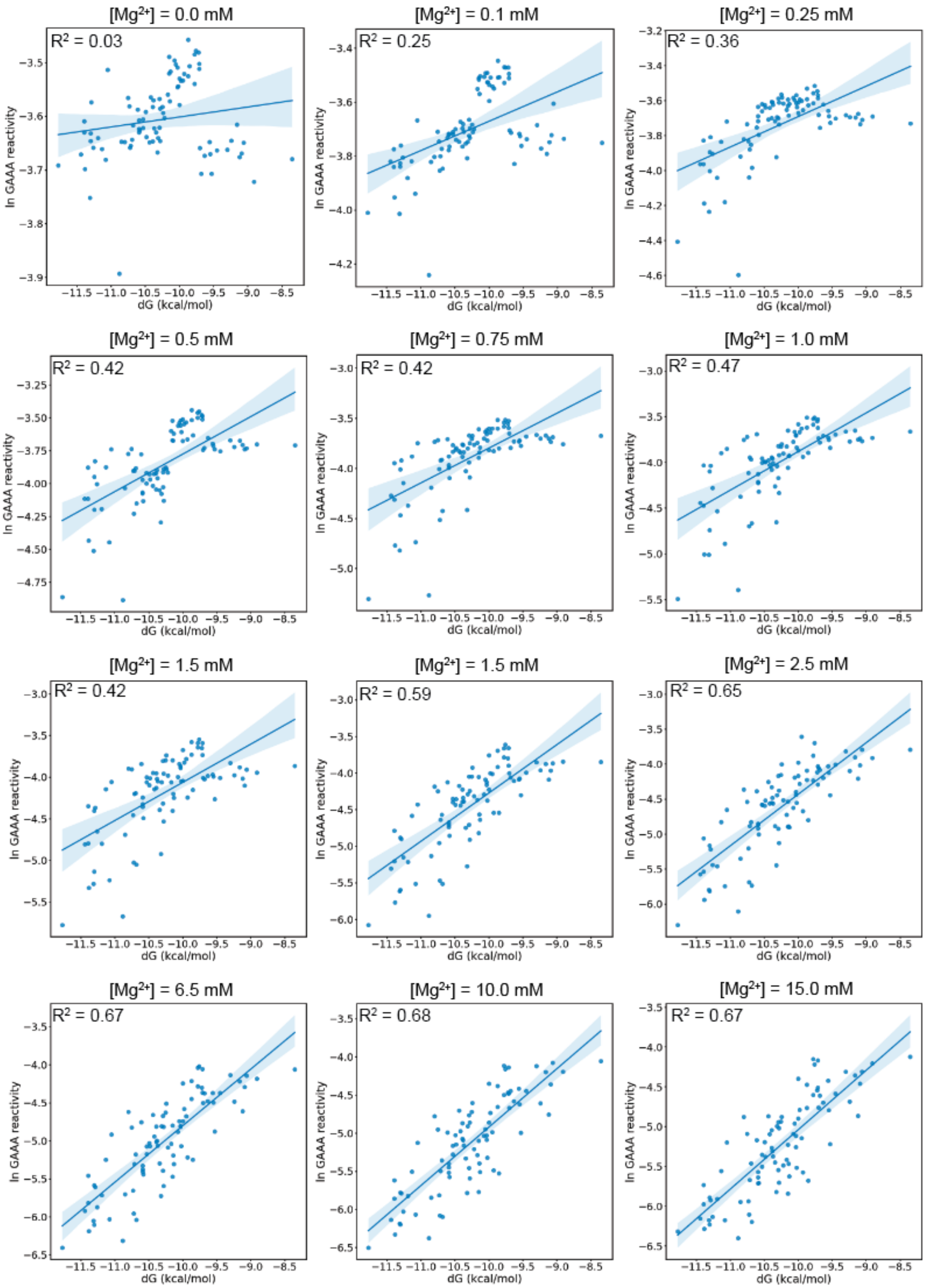
[Mg2+] = 0.0 mM
R2 = 0.03
[Mg2+] = 0.1 mM
R2 = 0.25
[Mg2+] = 0.25 mM
R2 = 0.36
[Mg2+] = 0.5 mM
R2 = 0.42
[Mg2+] = 0.75 mM
R2 = 0.42
[Mg2+] = 1.0 mM
R2 = 0.47
[Mg2+] = 1.5 mM
R2 = 0.42
[Mg2+] = 1.5 mM
R2 = 0.59
[Mg2+] = 2.5 mM
R2 = 0.65
[Mg2+] = 6.5 mM
R2 = 0.67
[Mg2+] = 10.0 mM
R2 = 0.68
[Mg2+] = 15.0 mM
R2 = 0.67
ln GAAA reactivity
dG (kcal/mol)

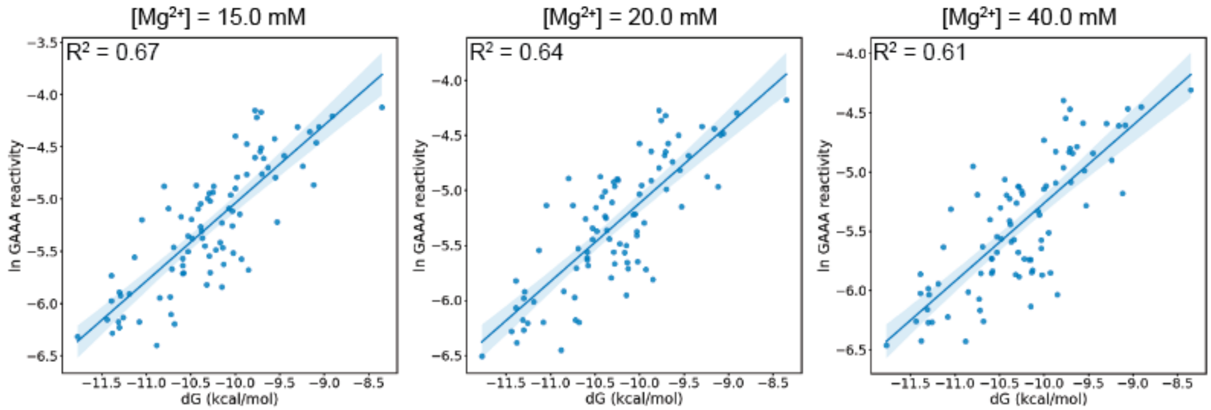

Supplemental Figure 14: Most $Mg^{2+}$ concentrations yield high correlations with RNA-MaP ΔG measurements.

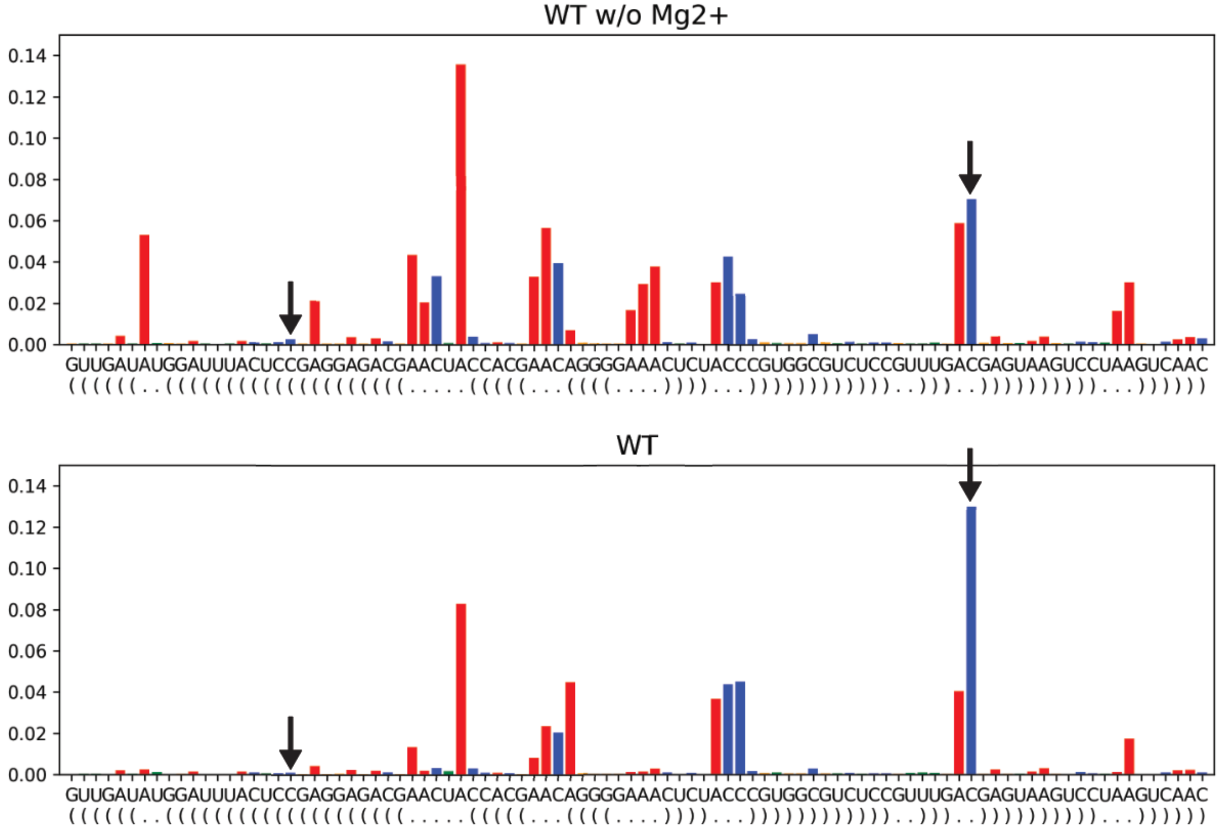

Supplemental Figure 15: Reactivity profile of kink turn of C-C pair like the C-C mismatch in the CCUAAC_CAUGG TLR variant.

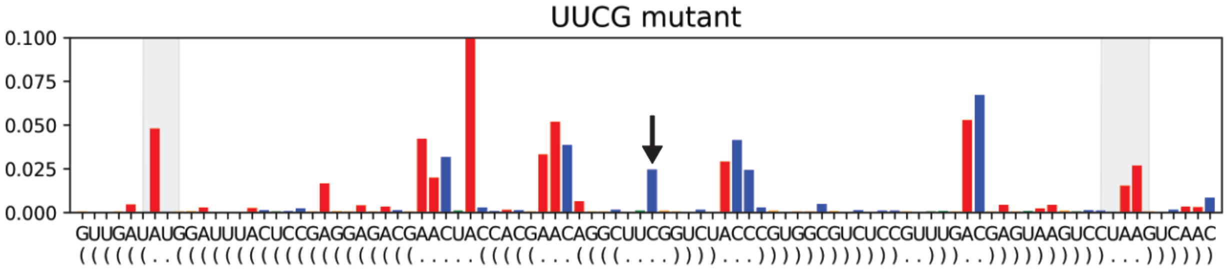

Supplemental Figure 16: reactivity average of C in UUCG, which is flipped out.

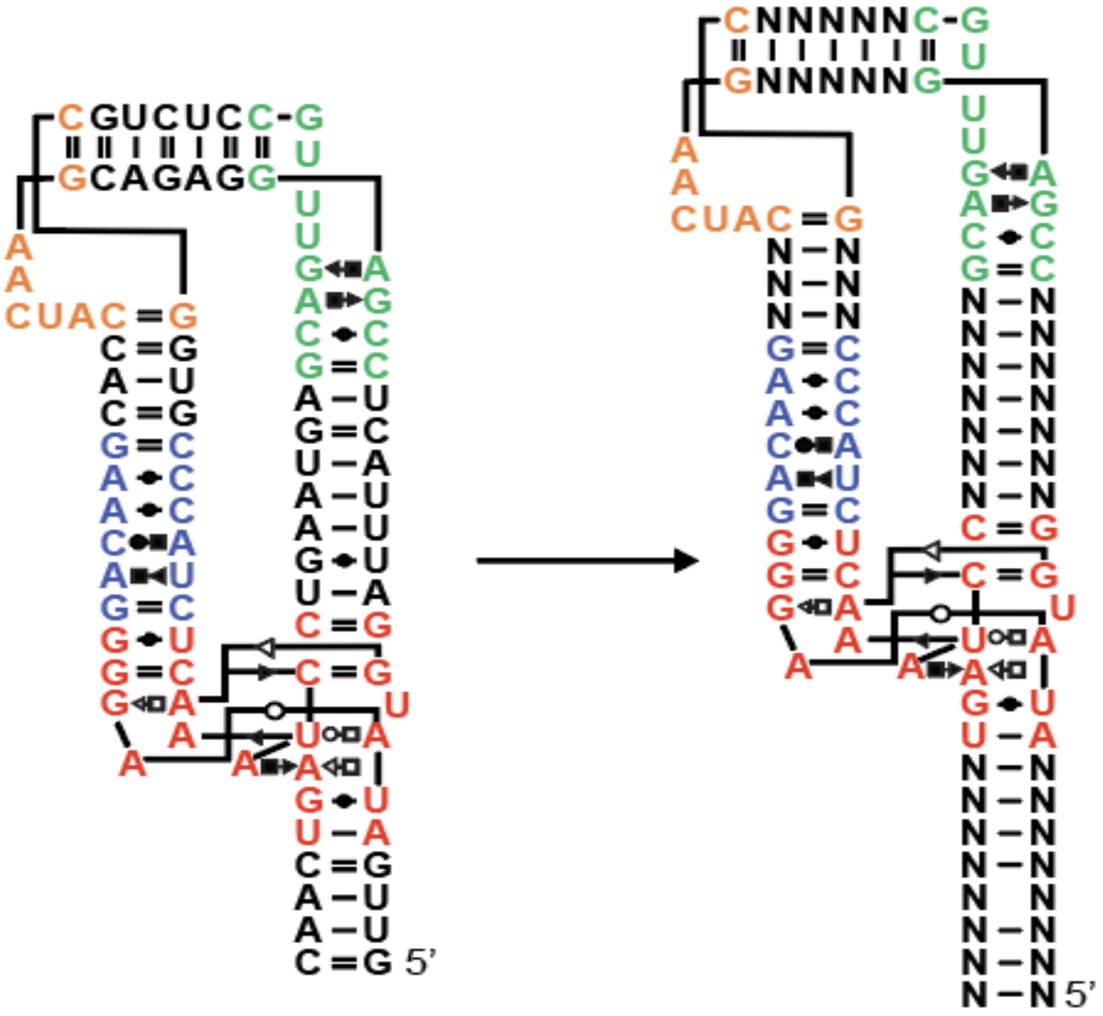

Supplemental Figure 17: Helix randomization strategy to increase diversity.
